# Discovery of the Honeycomb Synapse in Spinal Motor Circuits

**DOI:** 10.64898/2026.09.22.753133

**Authors:** Matthew J. Broadhead, Carlotta Löer, Janine Malseed, Katarina Parascandolo, Ani Ayvazian- Hancock, Anna N. Bak, Emma Butler, Molly Roberts, Simon A. Sharples, Leandro Lemgruber, Anthony Dornan, Jessica Valli, Bartosz Wasicki, Marcin Bączyk, Edita Bulovaite, Gábor Varga, Noboru H. Komiyama, Seth G.N. Grant, David I. Hughes, Claire F. Meehan, Gareth B. Miles

## Abstract

The structural and molecular diversity of synapses in the nervous system contributes to the specialisation of neural circuits underlying diverse behaviours. We have discovered a morphologically distinct postsynaptic specialisation in the mammalian spinal cord. We have named this the Honeycomb Synapse based on its elaborate postsynaptic nanostructure, comprising rings formed of ∼6 scaffolding protein domains that create multiple perforations throughout the large postsynaptic domain. Analysis of the organisation of other synaptic proteins reveals that Honeycomb Synapses harbour mixed signalling properties structurally facilitated by a postsynaptic scaffold matrix supporting chemical transmission, with gap junction proteins occupying some of the perforations. We reveal anatomical diversity in the presence of Honeycomb Synapses on populations of α-motoneurons across the lumbar spinal cord in mice from approximately 2 weeks of age through to adulthood. The Honeycomb Synapse was found to be a subtype of synapse within the Ia afferent monosynaptic stretch reflex circuit. Finally, we have identified that Honeycomb Synapses are highly vulnerable to degeneration in two different genetically engineered mouse models of Amyotrophic Lateral Sclerosis (ALS), contributing to monosynaptic stretch reflex circuit dysfunction. These findings suggest that synaptic diversity within circuits may confer selective vulnerability to distinct synaptic subclasses.

## Introduction

Synaptic diversity is critical for facilitating a breadth of resultant behaviours produced by the nervous system^1,2^. The diversity of synaptic function is determined in part by the molecular content of the synapse, such as the type of neurotransmitters released, the dynamic expression of different receptor subtypes and their organisation by different postsynaptic multi-protein complexes ^3–7^. Synaptic function can also be determined by the structure of the synapse. For example, the morphology of the dendritic architecture can modulate and gate molecular trafficking ^8,9^, and the number postsynaptic protein nanodomains within the synapse determines the strength of synaptic transmission ^6,10–13^. Therefore, synaptic diversity influences neural circuit output and the subsequent generation of myriads of behaviours.

Consider motor neurons (MNs) in the spinal cord and brainstem; the final common neuronal pathway to all our actions ^14^. These cells receive and integrate a wide range of synaptic inputs including monosynaptic inputs from sensory neurons, and a range of inhibitory, excitatory and modulatory inputs from local spinal interneurons and descending inputs from the cortex and brainstem ^15,16^. These connections are also subject to activity-dependent plasticity throughout early development and into adulthood ^17^. Understanding the structure, composition, and functional diversity of synapses onto MNs is likely to help advance our understanding of how these cells integrate diverse and generate complex motor patterns to influence behaviours.

Synapses are also highly vulnerable components of the nervous system in the early stages of many neurodegenerative disorders ^18–20^. In Amyotrophic Lateral Sclerosis (ALS), the most common motor neuron disease, there are pathologically conserved synaptic hallmarks including early-stage circuit hyperexcitability, maladaptive plasticity and synapse degeneration prior to MN loss ^20–23^. Identifying which types of synapses are vulnerable to pathological mechanisms could highlight the underlying mechanisms of the disease and reveal potential therapeutic targets ^24,25^.

In our ongoing research into the diversity of spinal cord synapses in health and disease, we serendipitously identified an unusual postsynaptic structure on MNs. These postsynaptic formations were distinguishable by their large size and numerous perforations within the postsynaptic matrix. Super-resolution microscopy revealed a complex nanostructural organisation of ∼6 postsynaptic nanoclusters of the scaffolding protein, PSD95, surrounding each perforation. Because of this distinctive morphology, we termed these structures Honeycomb Synapses. In this study, we aimed to define the structural and molecular properties of Honeycomb Synapses on to MNs, establish their prevalence throughout the lumbar spinal cord and during postnatal development, investigate their structural plasticity and also investigate their vulnerability in ALS.

## Results

### Initial Finding of Multiperforated Postsynaptic Densities

To investigate the diversity of excitatory synapse PSDs in the mammalian spinal cord, we examined the adult lumbar spinal cord sections of a genetically modified knock-in mouse model expressing eGFP-tagged postsynaptic scaffold protein, PSD95 (PSD95-eGFP ^26^) (Figure 1a). Using Airyscan microscopy, a high-resolution 3D Z-stack image was acquired from within the lateral ventral horn of the spinal cord where MNs reside (Figure 1b). From this acquisition, amongst thousands of PSD95 structures, a distinctive PSD was observed that was substantially larger than a typical PSD and comprised numerous holes or perforations (Figure 1c) (example is ∼4 μm in maximum length with over 10 perforations).

**Figure 1.**
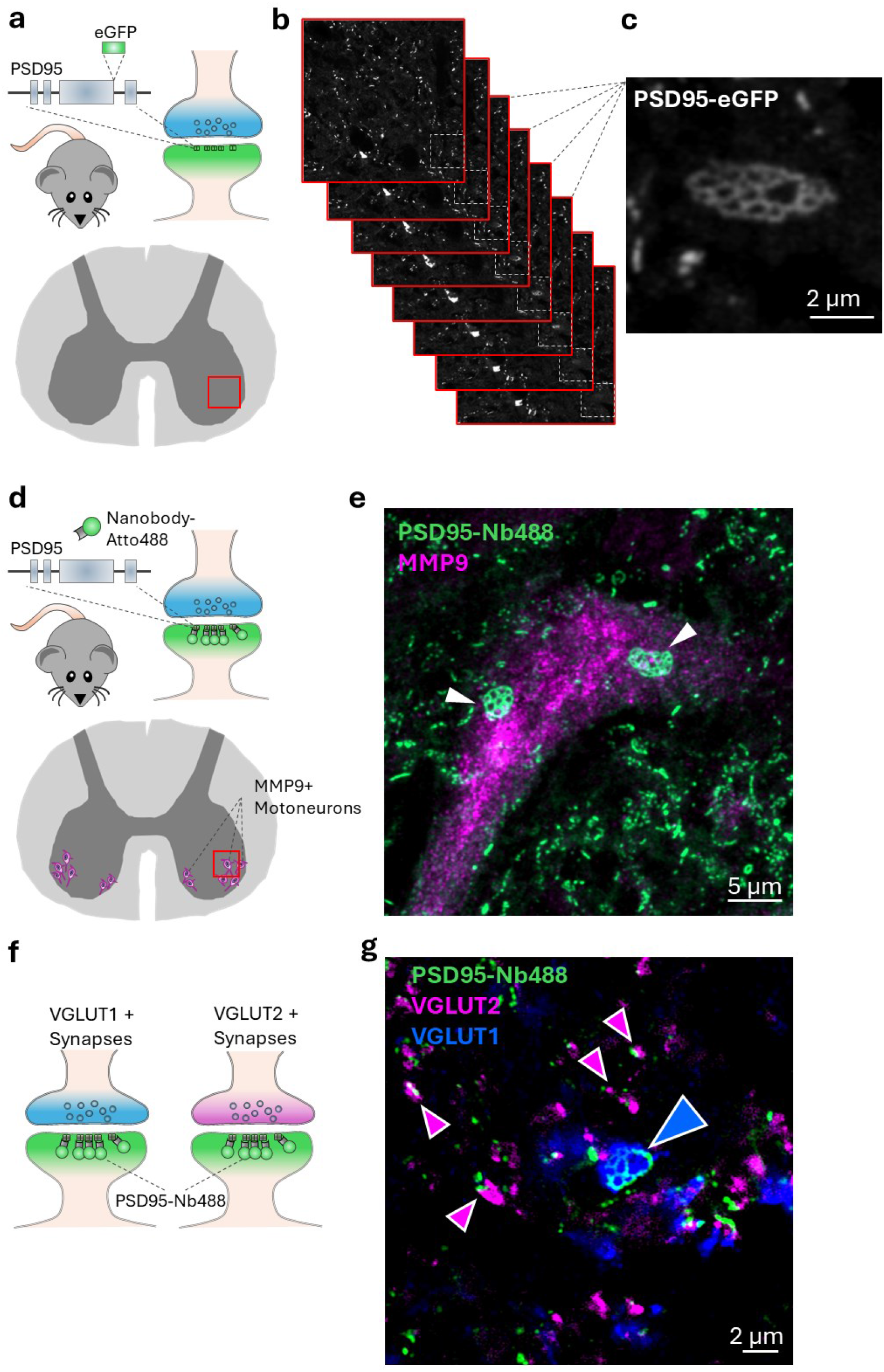
Discovery of a complex postsynaptic structure. a, Lumbar spinal cord tissue was obtained from an adult PSD95-eGFP homozygous mouse, in which the endogenous PSD95 (DLG4) is genetically tagged with eGFP to visualise postsynaptic scaffold proteins at excitatory synapses. Image was acquired from the lateral ventral horn (approximated by the red square). b, A 3D acquisition of PSD95-eGFP was captured with Airyscan microscopy, showing the sequential images through the tissue. A complex PSD95-eGFP structure was observed within the white box. c, Average projection of the stack reveals a congruent postsynaptic structure comprising a matrix of PSD95 interspersed with multiple perforations. d, PSD95 expression was visualised in the adult wildtype mouse spinal cord, immunolabelled using an anti-PSD95 Nanobody conjugated to Atto488 (PSD95-Nb488) and an antibody raised against MMP9 – a marker of fast-fatigable αMNs. e, Average-projection image from the Z-stack of an MMP9+ MN shows two example multiperforated PSDs onto the cell body of the neuron. f, Populations of excitatory synapses in the mammalian spinal cord can be labelled with presynaptic VGLUT1 and VGLUT2 in addition to postsynaptic PSD95. g, In an adult mouse spinal cord, a multiperforated postsynaptic structure is observed opposed to a VGLUT1 + presynaptic bouton (white/blue arrow), quite separate from VGLUT2 + synapses that show simpler morphologies (white/magenta arrows).

We next identified similar complex, multi-perforated PSDs, labelled with a fluorescently conjugated anti-PSD95 nanobody, on the cell body of an MMP9-labelled αMN (Figure 1d-e). These qualitative findings confirmed that the complex, multi-perforated PSD was a bona fide structure on spinal MNs that could be discerned through genetic and immunolabelling approaches. Furthermore, we identified that these multi-perforated PSDs were bona fide synapses that colocalised with the presynaptic bouton marker, VGLUT1 which contrasted to smaller non-perforated PSD95 clusters colocalised with smaller VGLUT2 labelled presynaptic boutons (Figure 1f-g).

These findings indicated that we had identified a rare structural form of excitatory synapse onto a population of MNs that could have a unique role in motor circuits. However, these initial qualitative findings were based on a small number of serendipitous observations. We therefore began to focus our investigations to better characterise and understand the nature of these novel synapses.

### Honeycomb PSD Nanostructure

To gain a more complete understanding of the structure of the multiperforated PSDs, we used different forms of high-resolution and super-resolution microscopy to visualise and characterise their morphology. Firstly, using Airyscan microscopy, we resolved that the postsynaptic domain was comprised of a ‘matrix’ of PSD95 interspersed with ‘perforations’ (Figure 1a). The expression of PSD95 within the PSD matrix is not homogeneous. PSD95 is known to form nanodomains or subclusters (herein termed ‘clusters’) with a diameter of ∼70-150 nm and the PSDs can be comprised of numerous clusters of PSD95^10,12,27^. Airyscan microscopy revealed this clustered organisation of PSD95 within the matrix of the multiperforated PSDs (Figure 2a), as indicated by the 5-6 peaks in fluorescence intensity around a perforation (Figure 2b).

**Figure 2.**
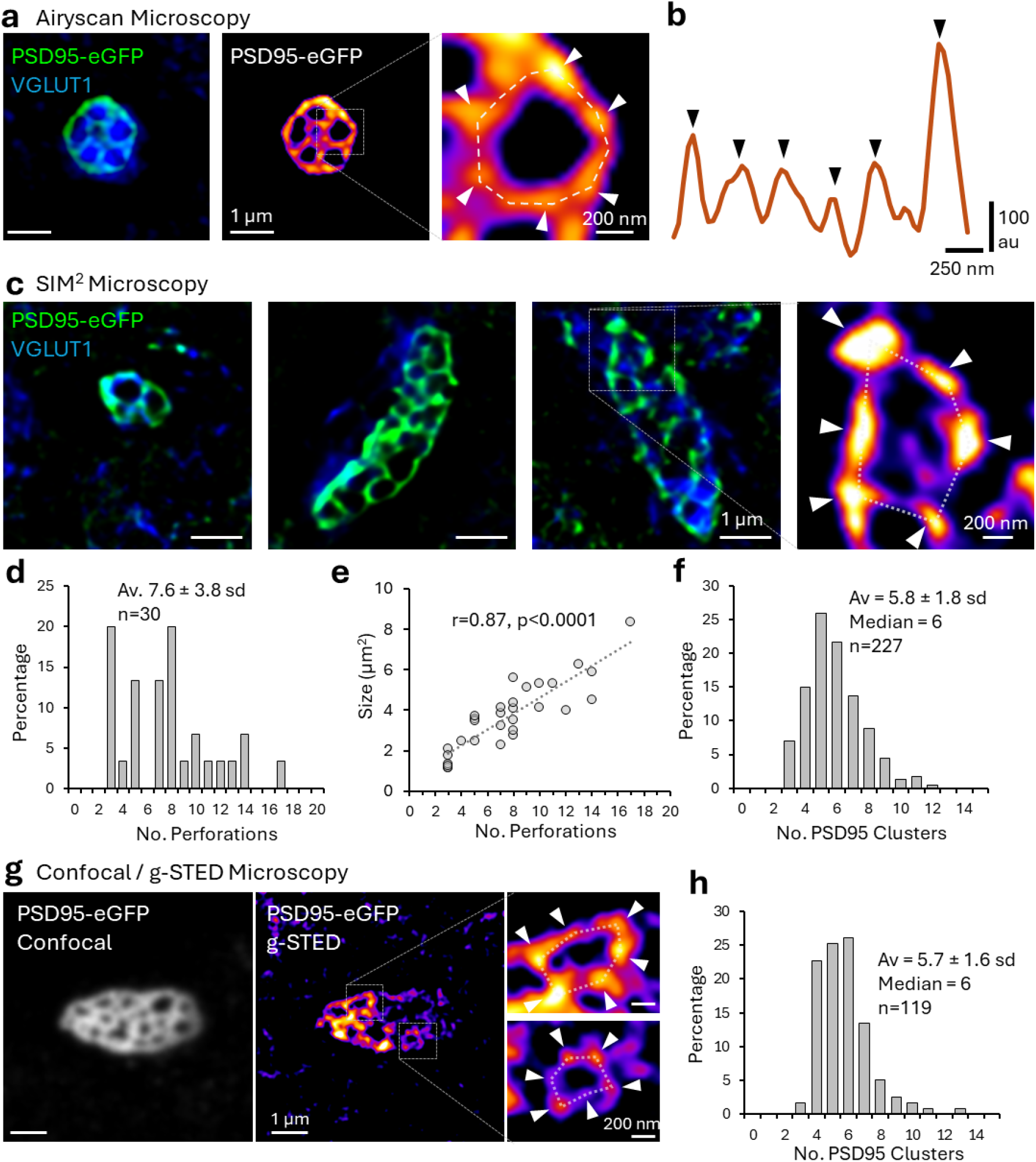
Super-Resolution Microscopy Reveals Honeycomb Nanostructure. a, Synapses in the adult mouse spinal cord expressing PSD95-eGFP, co-immunolabelled with VGLUT1, were visualised using Airyscan microscopy. Images display an average projection from across sequential Z-stacks to capture the whole synapse. Examining the PSD95-eGFP signal, clusters of PSD95 expression reside within the matrix of the synapse surrounding each perforation (white arrows). b, Intensity profile along the dotted line following the PSD95 clusters surrounding the perforation shown in panel (a) shows ∼6 peaks (black arrows), each ∼200 nm in diameter. c, SIM^2^ microscopy was used to identify and visualise these synapse types with a higher lateral and axial resolution in the adult mouse spinal cord expressing PSD95-eGFP, co-immunolabelled with VGLUT1. Example images of 3 multiperforated synapses are shown, and a cropped image (right) showing the organisation of 6 apparent PSD95 clusters (white arrows) around an individual perforation. d, histogram plots the range of the number of perforations per synapse, from a sample of n=30 synapses. Only VGLUT1+ synapses with 2 or more complete postsynaptic perforations were used for analysis. e, Scatter plot shows the positive correlation between the PSD size and the number of perforations. f, Histogram plots the distribution of the number of PSD95 clusters surrounding each perforation, from a sample of n=226 perforations across the n=30 synapses. g, Correlative confocal and g-STED microscopy was performed on synapses with multiple perforations in their PSD95-eGFP organisation. The number PSD95 clusters around each perforation (white arrows in the two cropped examples) was quantified. h, Histogram plots the distribution in the number of PSD95 clusters surrounding each perforation (n=119 perforations from a sample of 12 identified multiperforated synapses). From both the SIM^2^ and g-STED microscopy, the average number of PSD95 clusters surrounding each perforation was ∼6, therefore we henceforth term these Honeycomb Synapses.

To better resolve and quantify this organisation, we used super-resolution microscopy with more optimal resolution (sub-100 nm lateral resolution). Images of multiperforated PSDs (n=30) were captured using SIM^2^ and their substructure was analysed (Figure 2c). Multiperforated PSDs displayed an average of 7.6 perforations (ranging from 3-17) (Figure 2d). The size of the synapses averaged 3.7 µm^2^ (ranging from 1.1-8.3 μm^2^) and showed a significant, positive correlation with the number of perforations, indicating a modular organisation (Figure 2e). Analysis of the matrix of the PSDs revealed an average of 5.8 PSD95 clusters surrounding each perforation (median of 6 clusters; ranging from 3-12 clusters) (Figure 2f).

To validate these findings, we used gated stimulated emission depletion microscopy (g-STED) to visualise multiperforated PSDs (Figure 2g). Analysis revealed a similar nanostructural organisation to the postsynaptic matrix, comprising an average of 5.7 PSD95 clusters surrounding each perforation (median of 6 clusters, ranging from 3-13 clusters) (Figure 2h).

Taken together, nanoscopic analysis of these multiperforated PSDs reveals a complex, never-before-described organisation of groups of six postsynaptic scaffold protein clusters arranged circularly to form perforations within the large PSD domain. Due to its structure, we hereby defined these as Honeycomb Synapses.

### Molecular Organisation of the Honeycomb Synapse

To gain a more complete understanding of the molecular constituents of Honeycomb Synapses and their arrangement inside the synapse, we used a combination of genetically encoded fluorescence labelling and immunolabelling to detect synaptic proteins at Honeycomb Synapses, which were identified via colocalisation of PSD95eGFP and VGLUT1. Airyscan microscopy was used to acquire 3D images of Honeycomb Synapses that were then analysed from an average projection of the Z-stack. The degree of colocalization (Mander’s Coefficient) of different synaptic markers with either the PSD95 matrix or the perforations was analysed to assess their subsynaptic organisation (Figure 3a). The range of synaptic proteins that were assessed for expression within Honeycomb Synapses were categorised into 4 groups: Scaffolding Proteins (PSD93, n=8; SAP102, n=8; and Shank2, n=11; Supplementary Figure 1) Glutamatergic Receptors (NMDAR2A, n=9; GluA4, n=9; and GluA1, n=11; Supplementary Figure 2), Vesicular Release Site Proteins (Bassoon, n=10; Supplementary Figure 3) and Gap Junction Proteins (ZO-1, n=7; and Cx36, n=13; Supplementary Figure 3).

**Figure 3.**
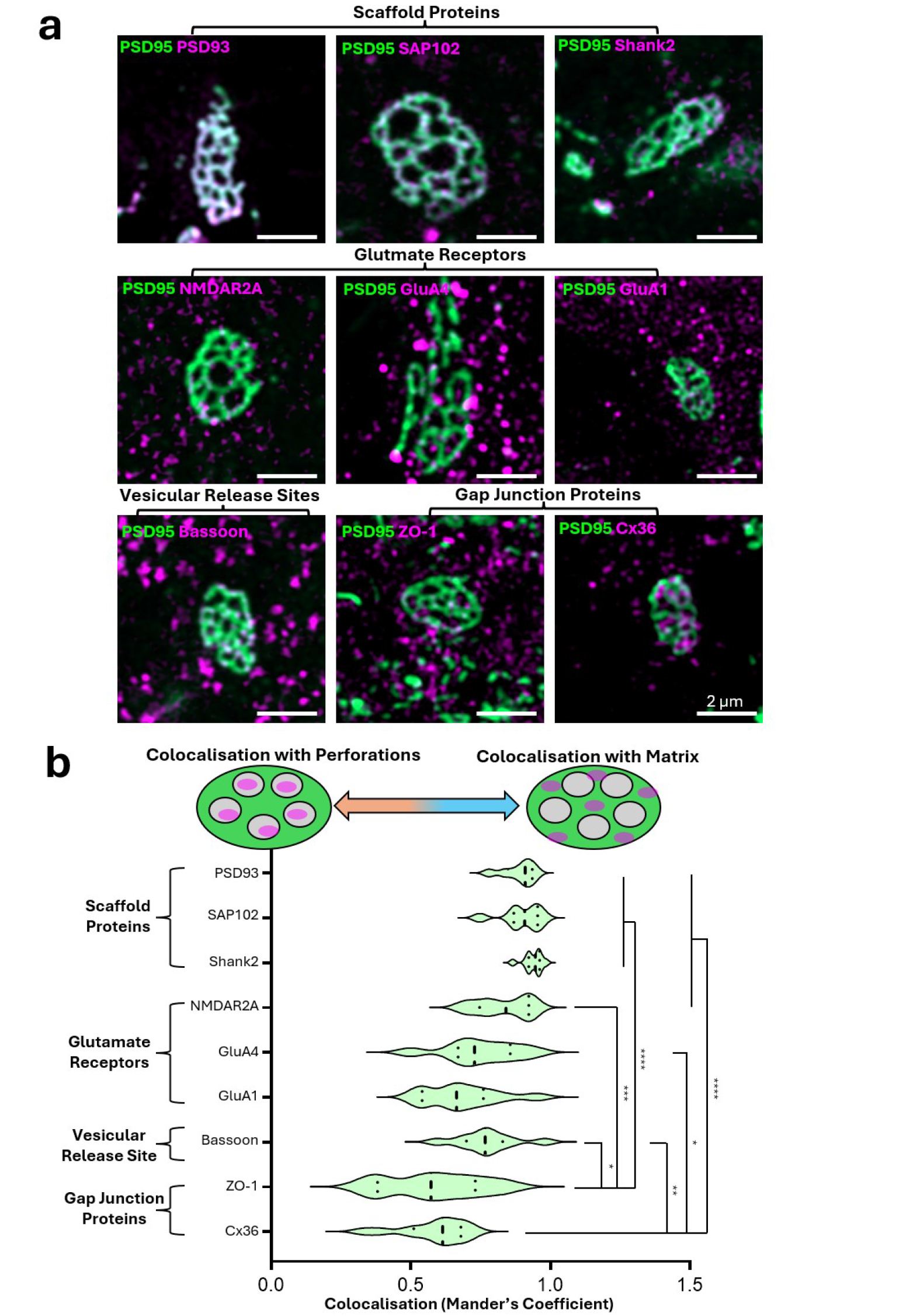
a, Images of Honeycomb Synapses (PSD95-eGFP) with other synaptic proteins, including Scaffold Proteins: PSD93 (n=8), SAP102 (n=8), Shank2 (n=11); Glutamate Receptors: NMDAR2A (n=9), GluA4 (n=9), GluA1 (n=11); Vesicular Release Protein: Bassoon (n=10); Gap Junction Proteins: ZO-1 (n=7), Cx36 (n=13). b, Bar chart plotting the degree of colocalization (Mander’s Coefficient) of each synaptic marker within the Honeycomb Synapse Matrix as discerned from PSD95-eGFP expression. Diagram above the graph represents how different proteins may be associated with the perforations or the matrix of the Honeycomb Synapse, depending on the Mander’s Coefficient score. One-Way ANOVA was performed (F(8,77)=14.7, p<0.0001), with a post-hoc Tukey’s multiple comparisons test. For clarity, only select pairwise comparisons are displayed.

We observed robust expression of PSD93 in Honeycomb Synapses, that colocalised with almost the entirety of the PSD95 matrix (Figure 3a; Supplementary Figure 1). We observed a lower expression of SAP102 within Honeycomb Synapses, forming smaller clusters that still aligned with the PSD95 matrix of the Honeycomb Synapses (Figure 3a; Supplementary Figure 1). Similarly, Shank2 formed fractured clusters that colocalised with the matrix of PSD95 (Figure 3a; Supplementary Figure 1). Therefore, the multiperforated appearance of the Honeycomb PSD was mostly visible by examining the expression of all four scaffolding proteins but was most clearly discerned with PSD95 and PSD93.

The glutamatergic receptor subunit proteins (NMDAR2A, GluA4 and GluA1) were all similarly expressed as small puncta localised within Honeycomb Synapses and positively associated with the PSD matrix, as expected based on their known interactions with PSD scaffold proteins (Figure 3a; Supplementary Figure 2). The presynaptic active zone marker, Bassoon, was presynaptically located but also aligned with the subclusters of PSD95 within the PSD matrix (Figure 3a; Supplementary Figure 2). This alignment of active zone – glutamate receptors – PSD scaffold subclusters is indicative of transsynaptic nanocolumns for glutamatergic transmission ^6^.

It was hypothesised that gap junctions may be expressed within the perforations of the Honeycomb Synapse based on prior evidence from perforated synapses within the crayfish nervous system ^28^. We used immunolabelling to visualise expression of Cx36 (the pore-forming component of gap junctions) and ZO-1 (a gap junction associated scaffold protein) at Honeycomb Synapses. We observed numerous clusters of Cx36 and ZO-1 at Honeycomb Synapses (Figure 3a; Supplementary Figure 3). Compared to the other synaptic proteins assessed, the punctate clusters of Cx36 and ZO-1 were more heterogeneously expressed throughout the postsynaptic domain, with some clusters aligning within the matrix, and notably other clusters aligning within the perforations of the Honeycomb Synapses (Figure 3a; Supplementary Figure 2).

Quantitative analysis of the colocalization (Manders Coefficient) of each tested protein with the PSD95 matrix of the Honeycomb Synapses revealed significant differences in the spatial organisation of Scaffolding Proteins, Glutamatergic Receptors, Vesicular Release Sites and Gap-Junction Proteins (Figure 3b; Supplementary Figure 4; F (8, 77) = 14.7, p<0.0001). The Scaffolding Proteins were significantly more colocalised with the matrix than the Gap Junction Proteins, ZO-1 and Cx36 (p<0.0001). Similarly, the glutamate receptor, NMDAR2A, was significantly more associated with the matrix than ZO-1 (p=0.0005) and Cx36 (p<0.0001). Bassoon was more significantly associated with the matrix than both ZO-1 (p=0.0175) and Cx36 (p=0.0026). To further validate these findings, we analysed the colocalization of each synaptic protein with the perforations of the Honeycomb Synapses and confirmed that the Gap Junction proteins were more significantly associated with the perforations than the other synaptic proteins analysed (Supplementary 5; F (8, 77) = 15.194, p<0.0001). It is also noteworthy that within each of the four groups of synaptic proteins (Scaffold Proteins, Glutamate Receptors, Vesicular Release Site, and Gap Junction Proteins) there were no significant differences in the colocalization of these proteins with PSD95, suggesting that each class of synaptic proteins displays a distinctive and discernible spatial organisation.

These data indicate that Honeycomb Synapses harbour a complex subsynaptic structure that consists of different subdomains for electrical and chemical signalling properties – with electrical transmission likely occurring within perforations and surrounded by an annulus of chemical transmission nanodomains.

### Honeycomb Synapses in Development and Ageing

We next asked when during development Honeycomb Synapses arise, and whether they persist into adulthood and during aging. Lumbar spinal cord tissue was obtained from PSD95-eGFP mice at postnatal day 3, 15, 30 and 360 of age (n=3 animals per time point). Tissue was immunolabelled with VGLUT1 and CHAT to visualise all the VGLUT1-PSD95 synapses onto MN somas (Figure 4a). Images were captured predominantly of lateral MNs in the ventral horn of upper lumbar spinal cord sections using spinning disk confocal microscopy. Semi-automated machine learning analysis within Imaris was trained and used to identify and classify the Honeycomb Synapses from within the pool of VGLUT1-PSD95 synapses.

**Figure 4.**
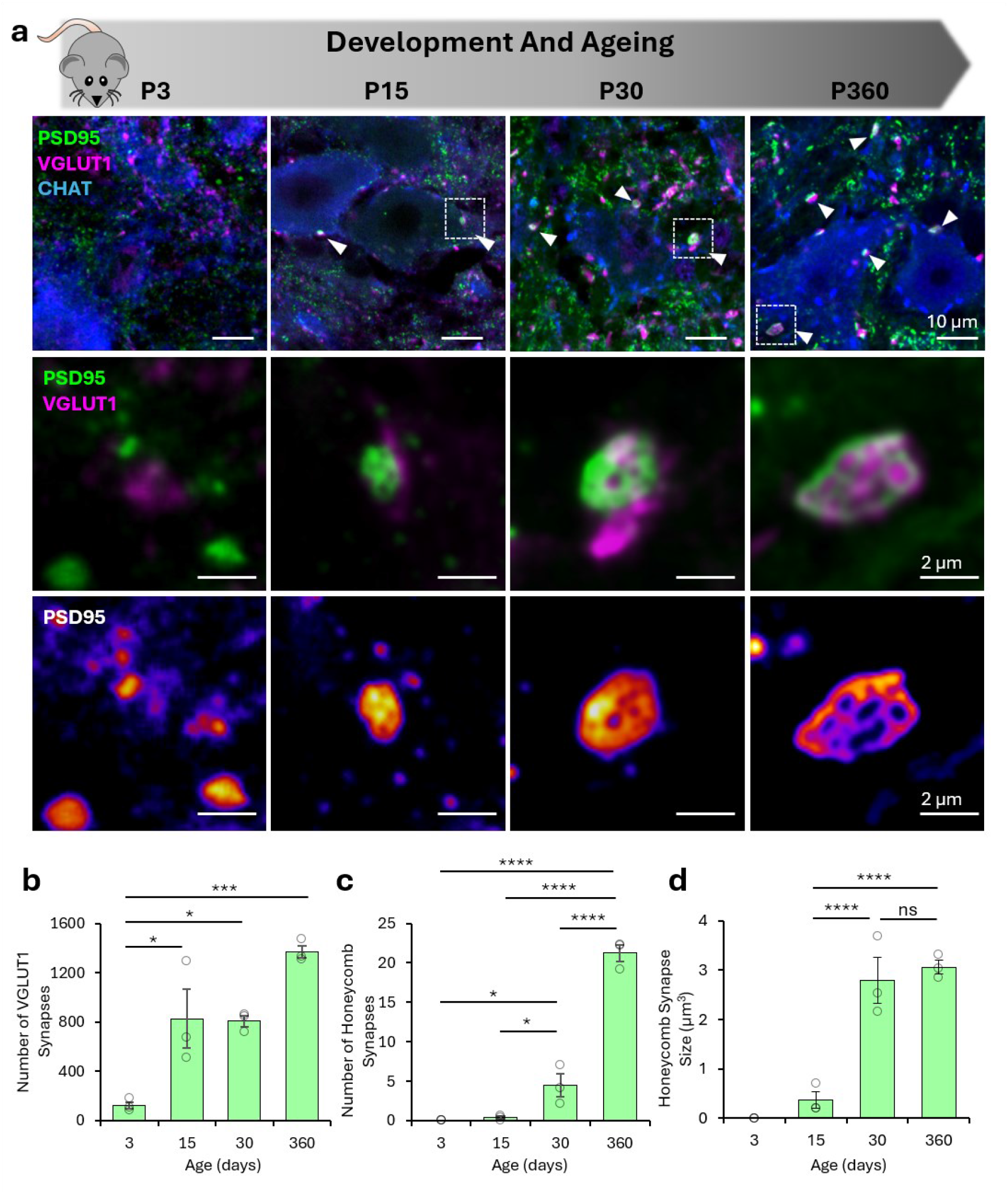
a, Analysis of Honeycomb Synapses across early development and ageing at 3, 15, 30 and 360 days old (n=3 per group). Tissue from PSD95-eGFP mice were immunolabelled with VGLUT1 and CHAT. White arrows denote individual Honeycomb Synapses. White boxes denote the cropped-out images below. b, Bar chart plotting the number of PSD95-VGLUT1 Synapses per image volume (221010.8 µm^3^) at each age. One-way ANOVA was performed (F (3, 8) = 16.79, p=0.0008), with a post-hoc Tukey’s multiple comparisons test. c, Bar chart plotting the number of Honeycomb Synapses per image volume at each developmental time point. One-way ANOVA was performed (F (3, 8) = 131.6, p<0.0001), with a post-hoc Tukey’s multiple comparisons test. d, Bar chart plotting the average size of Honeycomb Synapse PSDs in mice at each developmental time point. One-Way ANOVA was performed (F (3, 8) = 38.9, p<0.0001), with a post-hoc Tukey’s multiple comparisons test.

The number of all VGLUT1-PSD95 synapses onto MNs increased significantly across development (F(3,8)=16.8, p=0.0008, Figure 4b), increasing from 3 to 15 days (p=0.0168), remaining stable from 15 to 30 days (p=0.9996), and almost doubling between 30 and 360 days (p=0.051).

The number of Honeycomb Synapses classified from within the total pool of VGLUT1-PSD95 synapses also significantly increased over age (F(3,8)=131.6, p<0.0001, Figure 4c). No Honeycomb Synapse subtypes were identified in 3-day old animals, but a small number of Honeycomb Synapses were identified at 15 days. The number of Honeycomb Synapses then increased with significantly more identified at 30 days (p=0.042) and the greatest number observed at 360 days (p<0.0001). This developmental trajectory indicates that Honeycomb Synapses likely arise with the onset of weight-bearing motor behaviours, which occurs between the first and second week of age in mice, and that Honeycomb Synapse then continue to mature throughout adulthood. The structure of the Honeycomb Synapse PSDs changed significantly over age (F(3,8)=38.94, p<0.0001). Honeycomb synapse size increased between 15 and 30 days of age (p=0.0007) but did not change between 30 to 360 days (p=0.876), suggesting that their structure remains relatively stable into older age.

### Anatomical Mapping of Honeycomb Synapses

We next sought to understand whether Honeycomb Synapses were equally present on all lumbar spinal cord MNs, or whether there was diversity in Honeycomb Synapse number and structure that may be associated with functionally and anatomically distinct MNs and motor circuits. We therefore characterised the anatomical expression of Honeycomb Synapses within the mouse lumbar spinal cord. Quantifying synapses between the Lateral Motor Columns and Medial Motor Columns (LMC and MMC) would provide insights into whether Honeycomb Synapses were more predominantly associated with limb-based movements versus postural motor control respectively (Figure 5a). Meanwhile, analysis of synapses across lumbar segment 2 and lumbar segment 5 (L2 and L5) may provide insights into whether Honeycomb Synapses were associated more with flexor versus extensor motor units (Figure 5a). Spinal cord sections from 28-day old PSD95-eGFP mice (n=3) were obtained and immunolabelled with VGLUT1 and MMP9 to label the presynaptic boutons and αMNs. Images were acquired from the LMC and MMC of L2 and L5 spinal sections and Imaris was used for semi-automated classification and analysis of Honeycomb Synapses (Figure 5b).

**Figure 5.**
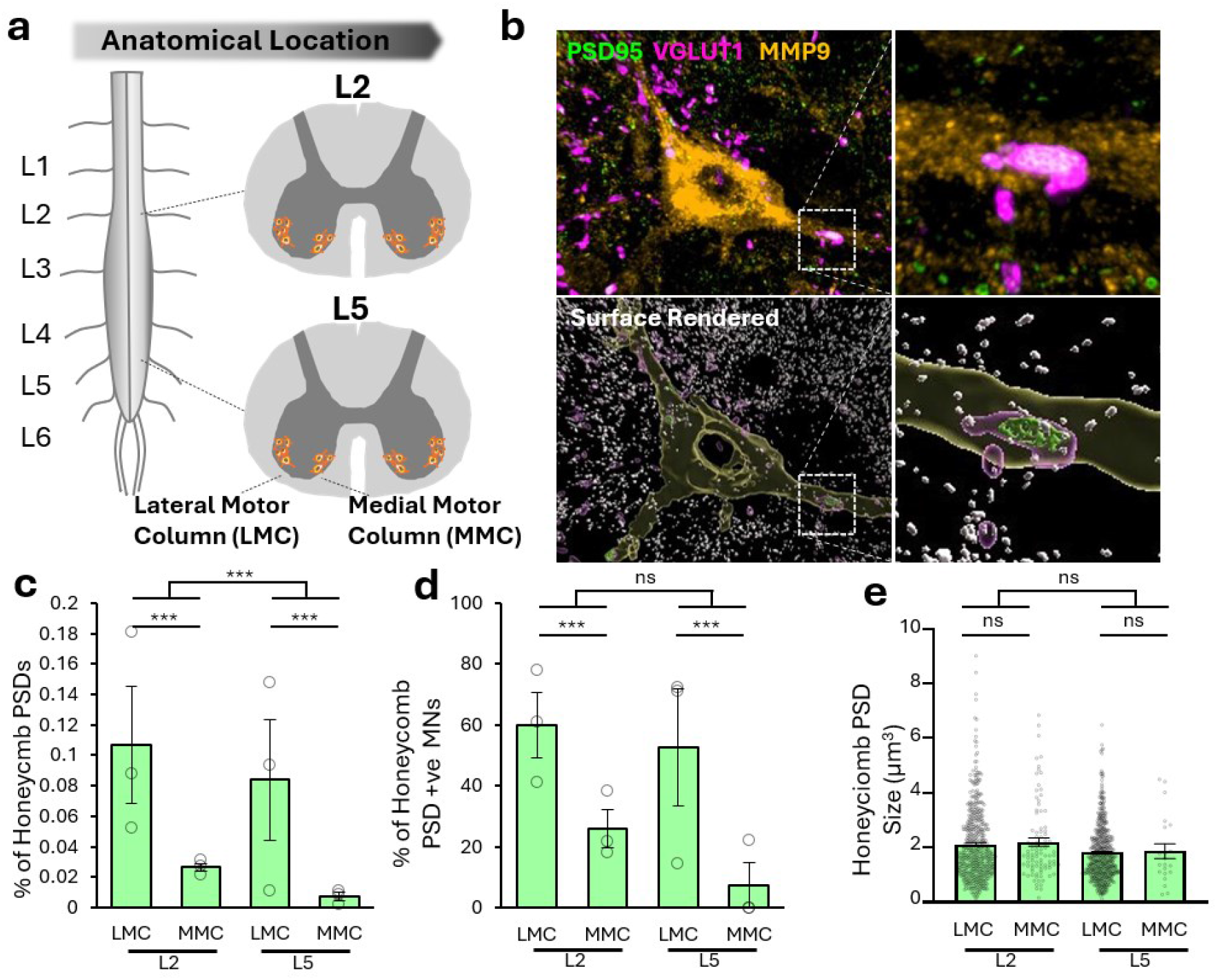
a, Diagram of the lumbar spinal cord of the mouse, illustrating the LMC and MMC at L2 and L5 where we surveyed for Honeycomb Synapses onto alpha-motoneurons (αMNs). b, Example of PSD95, VGLUT1 and MMP9 labelling with analysis classifying the Honeycomb Synapse. c, Bar chart plots the percentage of all VGLUT1-associated PSDs that were classified as Honeycomb Synapse PSDs. A generalised linear model (from n=3 animals) was used to analyse differences across the LMC and MMC in upper L2 and lower L5 lumbar segments. d, Bar chart plots the percentage of all MMP9-positive MNs that harbour at least 1 Honeycomb PSD across L2 and L5 in the LMC and MMC. e, Bar chart plots the size of Honeycomb Synapse PSDs across L2 and L5 in the LMC and MMC (across 1146 Honeycomb Synapses).

From analysis of over 2 million VGLUT1-associated PSD’s onto MMP9 neurons, Honeycomb Synapses were more frequently observed in the LMC compared to the MMC (β = 2.37, p < 0.001) and more frequently observed in L2 compared to L5 spinal cord segments (β = 0.97, p = 0.001) (Figure 5c). It was noted that ∼80% of the 1146 Honeycomb Synapses detected in this data set were associated with MMP9-labelled MNs, suggesting a predominant association with fast-fatigable αMNs. We next quantified the percentage of MNs that harboured at least 1 Honeycomb Synapse. Across the upper and lower LMC, 50-60% of MNs harboured Honeycomb Synapses whilst only 10-25% of MNs in the MMC harboured Honeycomb Synapses (β = 2.74, p = 0.001) (Figure 5d). There was no significant difference in the percentage of MNs containing a Honeycomb Synapse between L2 and L5 segments (β = 0.81, p = 0.479) (Figure 5d). The volume of MNs positively predicted the number of Honeycomb Synapses harboured on the neuron (Supplementary Figure 6a), suggesting that larger neurons were most likely to display Honeycomb Synapses. These findings indicate a preferential (but not exclusive) expression of the Honeycomb Synapse subtype on αMNs in the LMC of the upper lumbar spinal cord, associated with flexor motor units involved in limb movements.

Next, it was asked whether Honeycomb Synapse size was different between anatomical regions. Because some animals showed no Honeycomb Synapses in the MMC’s, we elected to pool analysis from the 1146 Honeycomb Synapses identified from across all 3 animals. There was no significant difference in Honeycomb Synapse size between the LMC and MMC (β = 0.286, p = 0.081) nor between L2 and L5 lumbar segments (β = 0.307, p = 0.097) (Figure 5e). Furthermore, MN volume did not predict the Honeycomb Synapse PSD volume (β = 0.00005, p = 0.102) (Supplementary Figure 6b). This would suggest that the size of the MN may determine whether they harbour Honeycomb Synapses but does not influence the structure of the Honeycomb Synapses.

### Functional Circuitry of the Honeycomb Synapse

We reasoned that the functional role of Honeycomb Synapses is to provide a specialised form of proprioceptive input to MNs, based on their close association with large VGLUT1-expressing boutons and their frequent occurrence within lateral motor pools containing large αMNs ^29,30^ (Figure 6a). To identify the functional circuitry of the Honeycomb Synapse, spinal cord tissue from Parvalbumin-Cre reporter line mice expressing YFP in Ia afferent sensory neurons was obtained and immunolabelled for PSD95 and VGLUT1 to identify the excitatory synapses and Honeycomb Synapses associated with Pv-Cre:YFP boutons (Figure 6b-c). 3D image acquisitions were acquired from the ventral horn of the lumbar spinal cord sections using high-resolution spinning disk confocal microscopy and Imaris was used for semi-automated classification and analysis of Honeycomb Synapses (Figure 6c).

**Figure 6.**
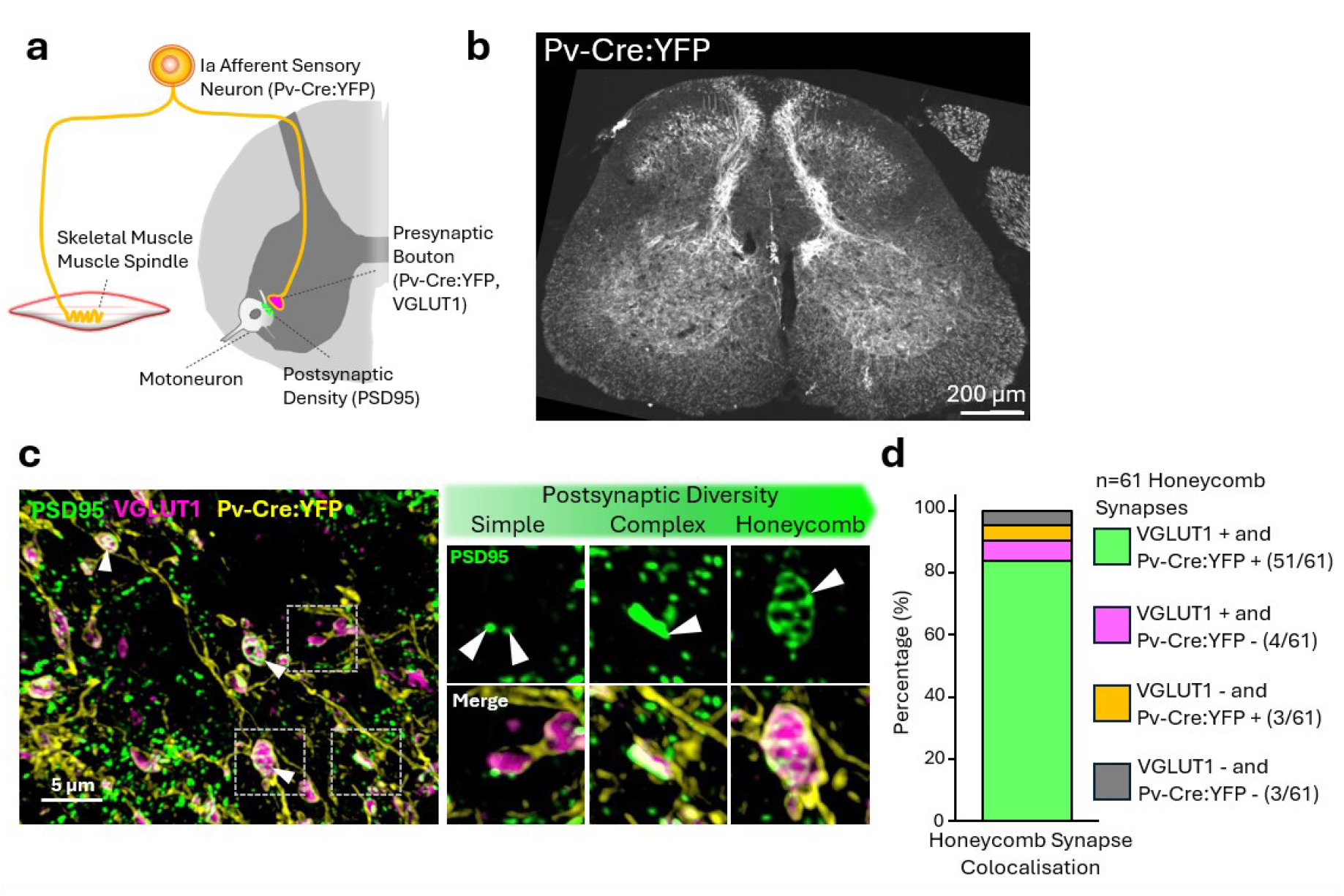
Honeycomb Synapses are part of the monosynaptic proprioceptive circuit to motoneurons. a, Diagram illustrating the monosynaptic proprioceptive circuit and the cellular/synaptic components for labelling using either genetic or immunolabelling approaches. b, Low magnification image of the lumbar spinal cord tissue from the Pv-Cre:YFP expressing mouse. c, Image displaying PSD95, VGLUT1 and Pv-Cre:YFP, with example synapses ranging in their perceived postsynaptic structural complexity, from Simple to Complex and ultimately Honeycomb structures. d, From a total of 61 Honeycomb Synapses identified from spinal cord sections across 2 mice, the percentages of these structures associated with either VGLUT1 and Pv-Cre:YFP (green), VGLUT1 alone (magenta), Pv-Cre:YFP alone (yellow) or neither (grey) are plotted, illustrating that a high proportion of Honeycomb Synapses are associated with the proprioceptive monosynaptic microcircuit.

From qualitative assessment, Ia afferent synapses (VGLUT1+/Pv-Cre:YFP+/PSD95+) were found to display a diversity of postsynaptic structures, from single, small PSDs to more structurally complex PSDs, and finally to those displaying multiple postsynaptic perforations consistent with Honeycomb Synapses (Figure 6c). From quantitative analysis of 61 Honeycomb Synapses identified from lumbar spinal cord sections across two Pv-Cre:YFP animals, 83.9% of the Honeycomb Synapses were associated with VGLUT1+/Pv-Cre:YFP+ presynaptic boutons. Meanwhile, 6.6% of Honeycomb Synapses were associated only with VGLUT1 and approximately 4.9% of Honeycomb Synapses were associated only Pv-Cre:YFP+ boutons (Figure 6d). These data indicate that Honeycomb Synapses are almost exclusively a structurally discernible subtype of the Ia afferent proprioceptive synapse. This result not only identifies the functional significance of this novel synapse for spinal cord motor control but also illustrates a considerable degree of heterogeneity within such a fundamental monosynaptic circuit

We did observe, however, that 4.9% of Honeycomb Synapses were not associated with either VGLUT1 or Pv-Cre:YFP boutons. From re-examination of our extensive analyses of VGLUT1-PSD95 synapses onto MMP9-labelled MNs in different anatomical subregions in the lumbar spinal cord (Figure 5), we confirmed that a similarly small population of non-VGLUT1-associated Honeycomb Synapses were discerned (Supplementary Figure 7a) that were significantly smaller in PSD volume than VGLUT1-associated Honeycomb Synapses (β = 0.746, p < 0.001, Supplementary Figure 7b). These findings indicate that a structurally smaller and much less frequent population of non-Ia afferent associated Honeycomb-Like Synapses are also present within spinal cord circuitry.

### Homeostatic Structural Plasticity in Honeycomb Synapse

Next, we wanted to investigate whether functional stimulation of this spinal circuit would evoke structural plasticity in the Honeycomb Synapses. To achieve this, we performed *in vivo* stimulation experiments. Adult wild-type mice underwent trans-spinal direct current stimulation (tsDCS); anodal stimulation was used to enhance Ia fibre activity, whilst cathodal stimulation was used to reduce Ia fibre activity compared to the sham control group ^31^. Stimulation was provided over 10 consecutive days before the animals were euthanised for perfusion fixation to isolate the lumbar spinal cord for immunohistochemistry. Honeycomb Synapses were visualised using high-resolution spinning disk confocal microscopy and analysed using semi-automated machine learning identification in Imaris. Honeycomb Synapses used for analysis were pooled from across multiple spinal cord sections from 2 sham animals (n=206 Honeycomb Synapses), 3 anodal stimulation animals (n=209 Honeycomb Synapses) and 2 cathodal stimulation animals (n=256 Honeycomb Synapses).

It was found that tsDCS stimulation evoked a significant effect on Honeycomb Synapse structure (*F*(2, 32819)=135.7, *p*<0.0001; Figure 7d). Whilst anodal stimulation did not induce significant structural alterations in Honeycomb Synapses compared to sham controls (p=0.350), cathodal stimulation resulted in significantly larger Honeycomb Synapses than sham controls (p<0.0001) and anodal stimulated (p<0.0001). This finding indicates a form of homeostatic structural plasticity, whereby the Honeycomb Synapse PSD area increases in response to chronically attenuated presynaptic input.

**Figure 7.**
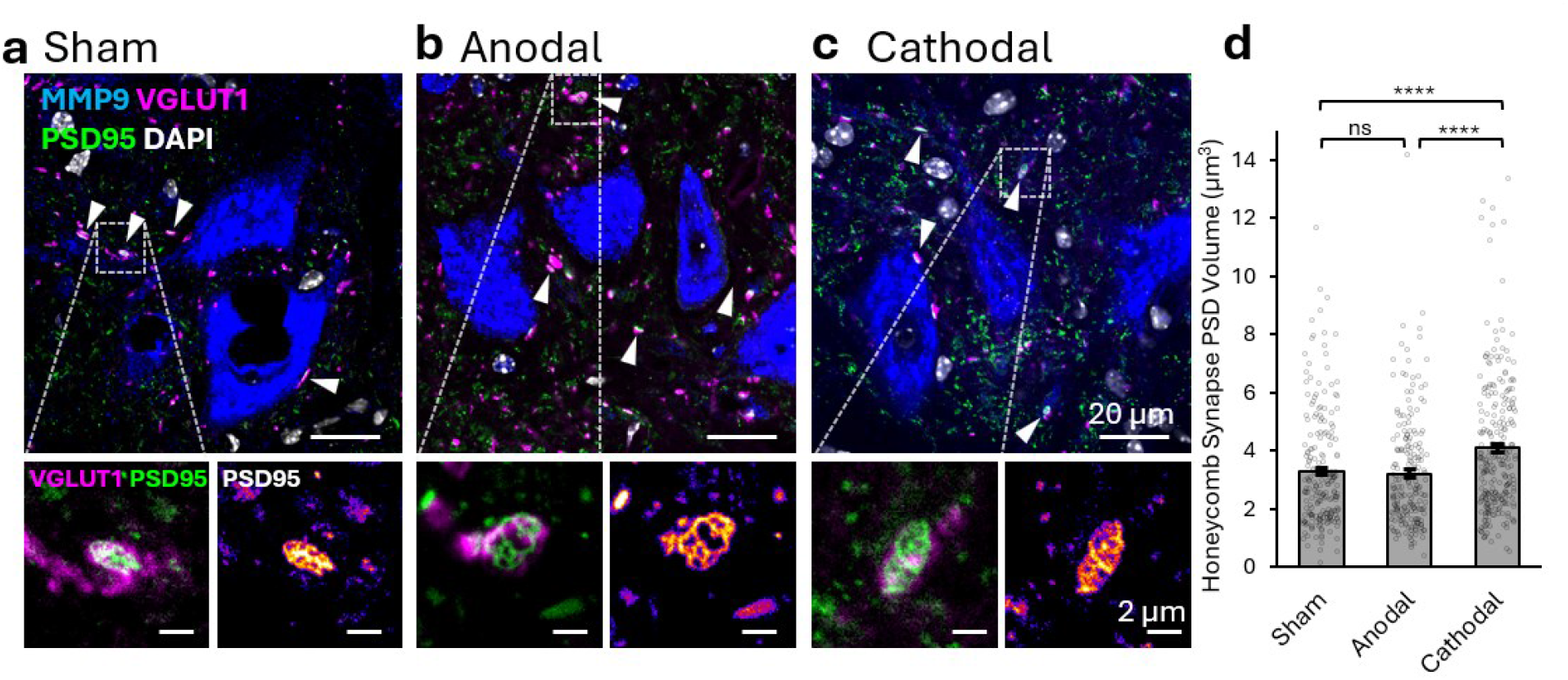
Honeycomb Synapse Structural Plasticity. High-resolution images were acquired of PSD95, VGLUT1, MMP9, and DAPI in the lumbar spinal cord of animals who underwent sham control tsDCS (a), anodal tsDCS (b) and cathodal tsDCS (c). White arrows indicate the presence of Honeycomb Synapses. Cropped images display example Honeycomb Synapse from each condition. d, Graph plotting the volume of Honeycomb Synapse PSDs. Data were analysed using a One-Way Welch ANOVA with Games-Howell post hoc tests for pairwise comparisons.

### Vulnerability of Honeycomb Synapses and Monosynaptic Reflex Circuit Dysfunction in ALS Models

Amyotrophic Lateral Sclerosis (ALS) is a fatal neurodegenerative disorder characterised by the progressive loss of MNs, leading to motor deficits, paralysis and ultimately respiratory failure ^22,32^. Synaptic pathology has been shown to arise prior to the loss of MNs ^20,33–35^. Changes in the Ia afferents to MNs have long been reported as a key hallmark of early synaptic dysfunction in ALS ^19,29,33,36–38^. We hypothesised that the integrity of large VGLUT1-associated Honeycomb Synapses may be most significantly impacted in ALS.

To address this hypothesis, we first utilised the SOD1^G93a^ (SOD1) mouse model, which recapitulates features of familial ALS including synaptic pathology, MN degeneration and reduced Ia afferent mediated H-reflex response ^33,39–41^. Spinal cord tissue was obtained from symptomatic SOD1 animals (n=7, 6 males, 1 female), and littermate controls (n=6, 3 males, 3 females) at a median age of 118 days and immunolabelled for PSD95, VGLUT1 and MMP9 (Figure 8a-b). The number of PSD95-VGLUT1

**Figure 8.**
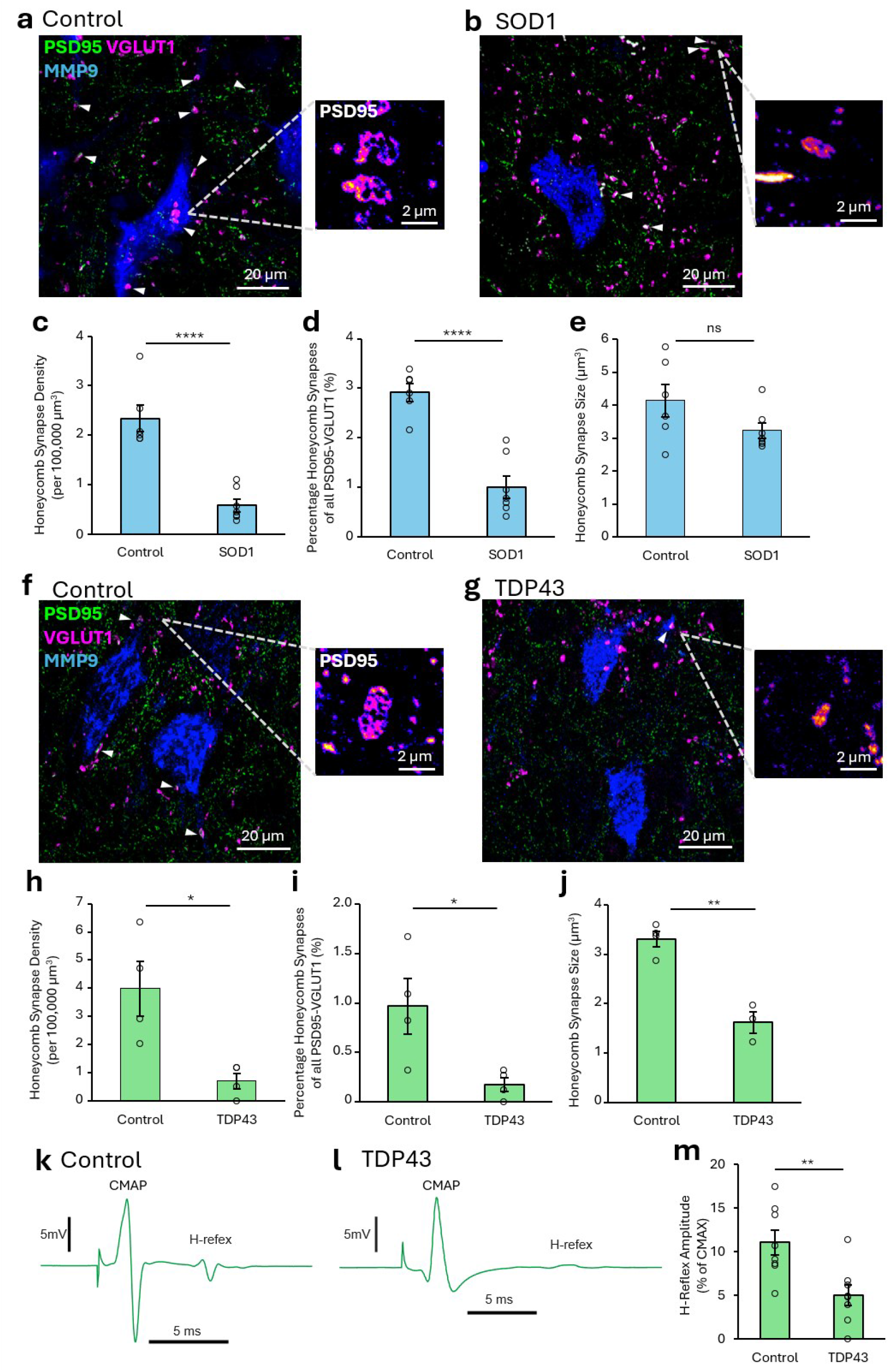
Honeycomb Synapse Degeneration and Circuit Dysfunction in Models of ALS. a-b, Images acquired of PSD95-eGFP, VGLUT1 and MMP9 in the lumbar spinal cord of control (a) and SOD1 (b) animals. White arrows highlight Honeycomb Synapses identified. The cropped images display example Honeycomb Synapses structures. n=6 control and n=7 SOD1 mice were used. Analysis was performed using a two-sample T-test. c, Graph plotting the number of Honeycomb Synapses in control and SOD1 mice. d, Graph plotting the percentage of all PSD95-VGLUT1 synapses that were classified as Honeycomb Synapses in control and SOD1 mice. e, Graph plotting the size of the Honeycomb Synapse PSDs in control and SOD1 mice. f-g, High-resolution images were acquired of PSD95, VGLUT1 and MMP9 in the lumbar spinal cord of control (f) and TDP43 (g) animals. White arrows highlight individual Honeycomb Synapses identified. The cropped images show example Honeycomb Synapse structures. n=4 mice per group. Analysis was performed using a two-sample T-test. h, Graph plotting the number of Honeycomb Synapses in control and TDP43 mice. i, Graph plotting the percentage of all PSD95-VGLUT1 synapses that were classified as Honeycomb Synapses in control and TDP43 mice. j, Graph plotting the size of Honeycomb Synapses in control and TDP43 mice. k-l, Example of averages of H-reflex recordings performed in control (k) and TDP43 (l) animals, 3-weeks post-induction. Example traces illustrate the compound muscle action potential (CMAP) and the subsequent H-reflex that was measured. m, Graph plotting the amplitude of the H-reflex (expressed as the % of the maximal CMAP – CMAX) in control and TDP43 mice. n=8 mice per group. Analysis performed using a two-sample T-test.

Honeycomb Synapses was significantly reduced by ∼75% in SOD1 mice compared to controls (t(11)=6.36, p<0.0001; Figure 8c). There was also a significant reduction in the percentage of PSD95-VGLUT1 synapses that were classified as Honeycomb Synapses in the SOD1 animals compared to controls (t(11)=6.56, p<0.0001; Figure 8d). Analysis of the sizes of the remaining Honeycomb Synapses in the SOD1 mice revealed no significant difference compared to those in the controls (t(11)=1.74, p=0.111, Figure 8e).

We then investigated Honeycomb Synapses in the inducible TDP43ΔNLS mouse model, which recapitulates features of sporadic ALS with TDP-43 mislocalisation, synaptic pathology and motor deficits ^19,42,43^. Bigenic animals were produced by crossing a tetO-hTDP-43-ΔNLS mouse with a NEFH-tTA mouse to produce progeny with tetracycline-repressible TDP-43 that lacks the nuclear localization signal (ΔNLS). When doxycycline is removed from the diet of bigenic animals, toxic mislocalisation of TDP-43 into the cytoplasm of neurons is induced ^42^. Between 2-4 weeks of induction, the TDP43 mice display TDP-43 mislocalisation and progressively severe motor deficits, both of which are absent in transgenic controls (expressing only NEFH-tTA) ^43^. To assess anatomically whether Honeycomb Synapses were impacted in the disease model, spinal cord tissue was obtained from 4-week induced TDP43 animals (n=4; 2 males and 2 females) and respective controls (n=4; 1 male and 3 females) and immunolabelled for PSD95, VGLUT1 and MMP9 (Figure 8f-g).

The number of PSD95-VGLUT1 Honeycomb Synapses was significantly reduced by ∼80% in TDP43 mice compared to controls (n=4 mice per condition; t(6)=3.27, p=0.0171; Figure 8h). Similarly, there was a significant reduction in the percentage of PSD95-VGLUT1 synapses that were classified as Honeycomb Synapses in the TDP43 animals compared to controls (t(6)=2.75, p=0.033; Figure 8i). In one of the TDP43 animals, no Honeycomb Synapses were identified at all. Analysis of the size of the remaining Honeycomb Synapse PSDs revealed that PSDs were significantly smaller in TDP43 animals compared to controls (t(5)=6.49, p=0.0013, Figure 8j).

To test the functional impact of this synaptic pathology, we measured the H-reflex (the electrical equivalent of the stretch reflex that is dependent on the Ia afferent synapse) in 3-week induced TDP43 mice and controls (n=8 per group; Figure 8k-l). H-reflexes were significantly reduced by ∼55% in the TDP-43 mice compared to controls (t(14)=3.26, p=0.0057; Figure 8m).

From across two distinct genetically engineered mouse models of ALS with different molecular mechanisms of the disease (recapitulating hallmarks of both familial and sporadic forms of ALS), we observe similar forms of synaptic pathology in the Ia-afferent associated Honeycomb Synapses. Namely, Honeycomb Synapses are significantly reduced by 75-80% in the diseased condition. These findings indicate that Honeycomb Synapse represent a particularly vulnerable subtype from within the broader pool of putative Ia afferent inputs to MNs. Furthermore, this loss of Honeycomb Synapses appears to correlate with significant reflex circuit dysfunction. Our findings further suggest that distinct disease mechanisms may differentially affect Honeycomb Synapse structure.

## Discussion

Synapse morphology and molecular organisation are key determinants of synaptic function and important features by which synapses from different circuits can be classified ^12,13,44,45^. There are many examples of specialised synaptic architectures, from the thorny excrescences of the CA3 region of the hippocampus ^12,46^, to the Calyx of Held in auditory processing circuits ^47–50^. Such unique synapses have provided models in which to investigate fundamental synaptic principles, whilst also offering novel insight into how specialised synapses help underpin the circuit output required to control different behaviours. Our discovery of the Honeycomb Synapse in spinal motor circuits may similarly provide new insights into sensory-motor physiology and advance our understanding of how synaptic organisation facilitates circuit function.

The spinal cord monosynaptic stretch reflex is one of the best characterised vertebrate neural circuits. Ia sensory afferents form excitatory glutamatergic synapses onto αMNs in the spinal cord to evoke a motor response in accordance with changes in muscle stretch ^29,51^. The electrically evoked signature of Ia afferent stimulation, the Hoffmann reflex (H-reflex), has become one of the most widely used experimental approaches for probing monosynaptic Ia afferent–αMN transmission, spinal excitability, and motor system function in both humans and animal models^52,53^. Together, the stretch reflex and H-reflex have been studied extensively for over 150 years, beginning with the first scientific descriptions of the reflex by Wilhelm Heinrich Erb and Carl Friedrich Otto Westphal in 1875 ^54^. Despite extensive study of this pathway for over a century, our findings reveal an unexpected level of synaptic specialisation within this canonical circuit.

Previous ultrastructural studies of Ia afferent synapses onto spinal MNs described presynaptic boutons with multiple active zones, most of which were small spherical or elongated structures, but 1-2% of which displayed a perforated appearance ^55^. Individual perforated active zones may colocalise with perforated PSDs, facilitating chemical transmission. Whilst we report a similar proportion of Honeycomb Synapses within Ia afferent synapses to these previously reported perforated active zone Ia afferent synapses, it is challenging to draw clear comparisons without direct correlative light-electron microscopy of the same synapses. Previous analyses using super-resolution microscopy confirmed that VGLUT1 synapses in the ventral horn are larger than VGLUT2 synapses and more frequently contact numerous, larger PSDs^27^.

Ultrastructural studies of synapses have long aimed to specify the functional significance of synapse constituents and their organisation in neural circuits and behaviours ^56,57^. Super-resolution microscopy reveals the complex organisation of synaptic proteins into either pre- or postsynaptic clusters or nanodomains. Larger synapses are built upon increasing numbers of these clusters, like conserved molecular building blocks ^10–12,44,58^. Honeycomb Synapses share some of the conserved structural organisation seen at other central glutamatergic synapses, with PSD95 and other scaffold proteins clustering neurotransmitter receptors and aligning with presynaptic vesicular release sites (Bassoon-positive clusters), forming transsynaptic nanocolumns within the synapse ^6^. Perforated PSDs have been described before in the mammalian brain from electron microscopy (EM) studies ^59^. In the dentate gyrus, perforated PSDs account for 16-25% of PSDs in young adult rats and it has been proposed these are intermediary stages in the dynamic cycle of synapse turnover ^59^. However, the scale of these perforated PSDs in the hippocampus is significantly smaller (∼400 nm diameter) than the Honeycomb Synapses we report here (1-5 µm diameter).

Analysis of the molecular components, and their subsynaptic organisation, within Honeycomb Synapses reveals a novel organisation facilitating mixed electrical-chemical signalling properties. There is considerable evidence from the literature that both spinal cord MNs and Ia afferents express Cx36 and utilise electrical transmission via gap junction complexes ^60–64^. What our findings now indicate is a specialised organisation of both chemical and electrical transmission machinery within the confined domains of a Honeycomb Synapse. Interestingly, this is consistent with prior evidence from the crayfish nervous system describing a mixed electrical-chemical synapse where an electrical synapse is surrounded by an “annulus” of vesicular-release machinery ^28^. Far from being passive conduits of current flow mediated only by connexin proteins, electrical synapses are increasingly recognised as complex structural signalling domains with a diverse molecular ensemble whose properties can dynamically shape transmission under physiological conditions ^65–67^. It is possible that the organisation of multiple chemical transmission sites surrounding each subsynaptic electrical transmission site contributes to the regulation of gap junction opening. Gap junction gating can occur through phosphorylation of Cx36, Ca²⁺-mediated calmodulin-dependent mechanisms, or other proposed biophysical processes, which may be facilitated at chemical synapses by the localized ion flux through glutamatergic receptors within these organised subdomains ^10,68^. This may be supported further at Honeycomb Synapses by the expression of GluA4 AMPARs in the PSD matrix, which exhibit fast kinetics and large Na+ conductance, permitting high fidelity transmission during high-frequency activity ^69,70^, and the expression of NMDAR2A within the PSD matrix, which facilitates high opening probability ^71^ and Ca^2+^ permeability ^72^. Taken together, we speculate that Honeycomb Synapses represent highly organised mixed electrical-chemical synapses that may be specialised to optimise the speed and reliability of signal transmission between Ia afferent terminals and MNs. Such an arrangement could be particularly advantageous during stretch reflexes, in which rapid and reliable recruitment of MNs is essential for the generation of an appropriate motor response.

From our analysis of the anatomical distribution of Honeycomb Synapses, we identified a form of diversity within a specific population of MNs based on whether they expressed Honeycomb Synapses. In the LMC, 50-60% of MMP9-labelled motoneurons harboured Honeycomb Synapses. MNs expressing high levels of MMP9 have been associated with large MN size, predominantly indicating high-threshold fast-fatigable αMNs ^73^. Assuming that MMP9-positive neurons constitute ∼50% of all lumbar MNs our findings suggest that a quarter of all MNs harbour Honeycomb Synapses. Why some MNs harbour Honeycomb Synapses, and others don’t, remains unknown – despite seemingly similar numbers of VGLUT1 inputs to such MMP9-labelled neurons. It is possible that Honeycomb Synapses are circuit-specific and derived from a particular type of sensory-motor unit. It is hypothesised that given Honeycomb Synapses were predominantly expressed on MMP9-labelled αMNs in the LMC, they may be associated with circuits innervating the fast fatigable muscle fibres of distal limb muscles ^74^. As these fast fatigable MNs have a higher threshold for recruitment, the electrical/chemical synaptic signalling properties of the synapse could facilitate this recruitment more rapidly and with greater force for the generation of vigorous reflex movements.

Related forms of synaptic diversity have also been observed at inhibitory synapses in spinal motor circuits. Differential organisation of Gephyrin, a scaffolding protein associated with glycine and GABA receptors, has been described between different spinal neurons ^75^. Renshaw cells display large, perforated rings of Gephyrin, termed ‘jaguar’ spots, which are associated with larger amplitude inhibitory postsynaptic currents. In contrast, MNs are innervated by inhibitory synaptic boutons associated with multiple small Gephyrin postsynaptic clusters, termed ‘cheetah’ spots, and these are associated more prominently with smaller amplitude glycine receptor-dependent postsynaptic currents ^75^. Together, these findings support the concept that postsynaptic nanoarchitecture contributes to the functional diversity among synapses both within and between spinal cord motor circuits.

Using tsDCS to functionally manipulate the activity of the spinal circuits *in vivo*, we observed a form of homeostatic structural plasticity to long-term suppression in pathway activity. Whilst the tsDCS technique may not be entirely selective to the Ia afferent pathway, it has been used previously to evoke changes in sensory-motor circuit physiology and as a means to investigate therapeutic strategies aimed at preserving these circuits against neurodegenerative disease mechanisms ^31,76^. The enhanced size of the Honeycomb Synapse PSD in response to chronic inhibition is consistent with pharmacologically induced nanostructural changes in excitatory synapse PSDs in response to tetrodotoxin blockade of neuronal activity *in vitro* ^10^, or the effect of astrocytic purine-mediated dampening of neuronal excitability in mammalian spinal cord ^27^. Homeostatic regulation of the post-synapse in response to chronic inhibition is associated with increased AMPA receptor content and an increased amplitude of postsynaptic currents ^77^. However, it was noted that Anodal (positive) stimulation of the pathway did not evoke a complementary reduction in Honeycomb Synapse postsynaptic size. We speculate that these large synaptic structures may be crucial for reliable fast synaptic transmission and may therefore be resistant to physiological mechanisms that would reduce their structural integrity. Nor did we observe any changes in the number of Honeycomb Synapses with either stimulation paradigm. Therefore, the complete mechanisms of synaptic plasticity within this circuit that regulate Honeycomb Synapse formation remain to be fully understood.

Synapses are highly vulnerable components of the nervous system in neurodegenerative conditions ^18^. Changes in Ia afferent synapses to motoneurons have been well documented in two of the most common forms of motor neuron disease, ALS and Spinal Muscular Atrophy (SMA) ^19,33,36,37,76,78,79^. Early-stage changes in inhibitory synapses have been suggested to play a key role in pathology in the SOD1 mouse model ^80,81^, whilst changes in other synaptic sources, such as local excitatory interneurons and modulatory cholinergic inputs to MNs, have been reported in mouse models and post-mortem tissue _19,20,33,43,82,83._

Analysis across the TDP43 and SOD1 models indicates highly conserved degeneration of Honeycomb Synapses by ∼80%. Both animal models also show, not only a loss, but a reduced proportion of Honeycomb Synapses within the entire class of VGLUT1 afferents to MNs. This indicates that ‘synaptic diversity’ itself is reduced in ALS. Our reflex studies in the TDP43 mice confirm that Ia afferents are less effective at bringing MNs to threshold, as has been described previously in the SOD1 model ^39^. Whether the reduced amplitude of the H-reflex is caused by the loss of Ia afferents or functional impairment of the remaining Ia afferents, remains to be understood. Nevertheless, the ∼55% reduction in the strength of the H-reflex is consistent with anatomical results, both with respects to Honeycomb Synapse loss reported here, and VGLUT1-associated synapse loss reported previously ^19,37^.

It is not clear why the TDP43 model displayed both a loss of Honeycomb Synapses and structural changes in the remaining synapses, whereas the SOD1 model only displayed synapse loss. Choosing comparable time points between two different genetic mouse models of ALS is challenging. The phenotypic severity of the inducible TDP43 model at this stage may be greater than that of 4-month-old SOD1 mice ^42^. However, TDP-43 has known roles in the translation of synaptic proteins such as GluA1 and is frequently localised within VGLUT1-expressing spinal cord synapses, which may contribute to primary pathologies at Honeycomb Synapses ^37,84^. Distinct pathogenic mechanisms may thus produce overlapping, but mechanistically separable, forms of synaptic pathology. Investigating a broad range of ALS models may therefore help distinguish conserved disease hallmarks from subtype-specific alterations and improve stratification of a disorder already highly heterogeneous in its genetic basis, clinical presentation, and progression. As we hypothesise that Honeycomb Synapses may be required for fast, high-fidelity glutamatergic transmission, it is therefore highly plausible that these connections may be more vulnerable than most to glutamate-mediated excitotoxicity given the evidence of early-stage hyperexcitability in spinal neural networks in ALS ^38,85^. Investigating the means to pharmacologically, genetically or functionally manipulate and preserve these specific Honeycomb Synapses may be a valid approach toward treating some of the underlying progressive motor deficits _in ALS 24,25,76,86._

In summary, our research reveals a never-before-described synaptic morphology that forms a distinct sub-class of sensory-motor circuit synapse onto αMNs in the spinal cord. The unique structural and molecular organisation of the Honeycomb Synapse is likely to confer distinct, as yet undefined, functional properties on Ia afferent input to MNs. The selective vulnerability of these synapses in murine models of ALS serves to reflect their importance in normal motor function. Understanding the precise function of Honeycomb Synapses will require complex interrogation to experimentally record from, or manipulate, individual Ia afferent synapse subtypes.

## Author Contributions

MJB conceptualised and directed the research, acquired funding for the research, performed experiments, analysis, interpretation of research findings and wrote the manuscript. CL, JM and KP performed experiments, analysis, interpretation of research findings and writing of the manuscript. AA-H, EB (St Andrews) and MR contributed to experiments. SAS provided conceptual and intellectual input and revisions of the manuscript. DIH provided tissue resources, provided conceptual and intellectual input and provided revisions of the manuscript. LL, AD and JV provided essential technical support conducting super-resolution microscopy experiments and were involved in the acquisition of grant funding to directly support the research. ANB and CFM provided tissue resources, conducted electrophysiology experiments of CMAP recordings, provided conceptual and intellectual input and revisions for the manuscript. BW and MB conducted in vivo experiments, provided tissue resources and provided revisions of the manuscript. EB (Edinburgh), GV and NHK were involved in the design and generation of the DLG-knock-in mouse models for visualising multiple scaffold proteins and provided tissue resources. SGNG acquired funding for the generation of the used DLG knock in mouse models, provided conceptual and intellectual input and provided revisions of the manuscript. GBM acquired funding for the research, provided conceptual and intellectual input and provided revisions of the manuscript.

## Acknowledgements

We would like to acknowledge the various members of the “Spinal Cord and Movement” and “Neural Circuits and Behaviour” groups at the University of St Andrews as well as the “Scottish Microscopy Society” and “Motor Control: Spinal Circuits and Beyond (St Andrews)” symposiums. Together, these communities of neuroscientists and microscopists have helped provide forums for academic discussion and support throughout the course of this research. For the purpose of open access, the author has applied a CC-BY public copyright licence to any Author Accepted Manuscript version arising from this submission.

## Data Availability

The research data supporting this publication will be made freely available in an open access data repository following acceptance to a peer-reviewed journal.

## Methods

### Animal Models and Ethics

Procedures performed on animals in the UK were conducted in accordance with the UK Animals (Scientific Procedures) Act 1986. Procedures conducted at the University of St Andrews were approved by the University of St Andrews Animal Welfare and Ethics Committee. Procedures performed at the University of Glasgow were approved by the Ethical Review Process Applications Panel of the University of Glasgow. Procedures performed at the University of Edinburgh were approved by The Edinburgh University Director of Biological Services. Procedures performed at the Poznań University were approved by the Local Ethical Committee (approval number 44/2018; Poznań, Poland). Procedures performed on animals in Denmark were conducted in accordance with the EU Directive 2010/63/EU and approved by the Danish Animal Experiments Inspectorate (Permission number 2022-15-0201-01327).

Tissues sourced from genetically engineered mouse models used in this study included: C57BL/6J (Charles River), PSD95-eGFP ^26^, SAP102-mKO2 ^26^, Ai32:B6;129S-Gt(ROSA)26Sortm32(CAG-

COP4*H134R/EYFP)Hze/J (Parvalbumin Cre – YFP), B6SJL-TgN(SOD1-G93A)1Gur/J (SOD1), and TDP43ΔNLS (TDP43) tissue obtained from the cross of tetO-hTDP-43-ΔNLS line 4 (https://www.jax.org/strain/014650) with a NEFH-tTA line 8 (https://www.jax.org/strain/025397). PSD93-HaloTag mice were generated as described ^87,88^. The genetic targeting strategy was used to fuse the HaloTag protein to the C-terminus of endogenous PSD93. HaloTag coding sequence (Promega) together with a short linker were inserted into the open reading frame of the mouse *Psd93* gene (DLG2). To establish triple-transgenic mice for visualising all three postsynaptic scaffolding proteins (PSD95, SAP102, PSD93), double homozygous PSD95^eGFP/eGFP^;SAP102^mKO2/mKO2^ mice were crossed with PSD93^HaloTag/HaloTag^ mice. SiR-Halo ligand was dissolved in DMSO to a stock concentration of 5 mM. The HaloTag ligand solution for injection was prepared at a concentration of 1.5 mM, consisting of 60 μl of 5 mM HaloTag stock solution, 20 μl of Pluronic F-127, and 120 μl of saline. A total volume of 200 μl of the HaloTag ligand solution was injected into each animal, as described previously ^87^. Prior to injection, animals were placed in a heat box for 5–10 minutes to allow the blood vessels of the tail to dilate and become more visible. The mice were placed in a rodent restrainer for the injection. A bolus injection of HaloTag ligand solution was performed into the lateral tail vein. Following injection, the animals were monitored for any adverse effects twice daily for the length of the experiment.

### Trans-Spinal Direct Current Stimulation (tsDCS)

The tsDCS procedures were conducted under isoflurane anaesthesia similarly as in ^89^, and the electrode arrangement and current intensity were adjusted as described in ^90^ and ^91^. Briefly, adult male C57BL/6J mice were assigned to sham, anodal, or cathodal tsDCS groups. To ensure a proper contact between the electrodes and the skin, animals’ backs were shaved prior to the tsDCS application. First, a tsDCS-type determining stainless steel rectangular electrode (2×3 mm), was positioned on the back of the animal over the Th10 vertebra, while the passive electrode (4×6 mm) was positioned over the sacrum ± 21 mm caudally. Direct current (60 µA) was applied for 15 min, with the rostral electrode serving as the positive electrode for anodal stimulation and the negative electrode for cathodal stimulation. Experimental stimulation was repeated for 10 consecutive days. Sham stimulation consisted of identical electrode placement and experimental handling, without the application of current.

### H-reflex Tests

TDP43 mice (n = 8) and control mice (n = 8; tetracycline activator-only mice) that underwent 3 weeks of doxycycline withdrawal to induce pathology were anaesthetised with Hypnorm (fentanyl citrate, 0.315 mg/mL, and fluanisone, 10 mg/mL) and midazolam (5 mg/mL), diluted 1:1 in distilled water. Custom-made wire needle electrodes were inserted into the intrinsic plantar foot muscles, and the medial plantar nerve at the ankle was stimulated at intensities sufficient to elicit a maximal CMAP (C-MAX). Signals were amplified using custom-made amplifiers (University of Copenhagen), digitised using a Power 1401 (CED, UK), and recorded, averaged, and analysed using Signal software (CED, UK).

The stimulation intensity was then reduced to determine the maximal H-reflex, and its amplitude was measured from averages of multiple trials and expressed as a percentage of C-MAX.

### Perfusion and Tissue Collection

Unless otherwise specified, tissue was collected from animals by first performing euthanasia with sodium pentobarbital (120 mg/kg) followed by transcardial perfusion with Dulbecco’s Phosphate Buffered Saline (D-PBS, Ca^2+^ - and Mg^2+^ -free; ThermoFisher Scientific #14190136) then 4% paraformaldehyde (PFA; ThermoFisher Scientific #053368.9M). For adult mice, 10 ml D-PBS and 10 ml PFA were used to perfuse each animal. For 15-day-old animals, the volumes were reduced to 7.5 ml, and for 3-day-old animals the volumes were reduced to 5 ml. Spinal cords were then removed and post-fixed for 2-4 h in 4% PFA. Spinal cords were then immersed in 30% sucrose in phosphate-buffered saline (PBS) for 48-72 hours before cryo-embedding in OCT compound and long-term storage at −80 °C. Cryosections at 20-30 μm thickness were obtained from lumbar spinal cords using a cryostat (either a Leica CM1860 or Epredia NX50), mounted on histology slides (Leica XTRA Adhesive slides, 3800200ae; or Leica APEX Adhesive slides, 3800080E) and stored at −80 °C for long-term preservation. For investigations into the molecular components of synapses, PSD95-eGFP animals that were immunolabelled with either Shank2, NMDAR2A, GluA4, GluA1, Bassoon, Cx36 or ZO-1 were perfused with 1% PFA due to evidence that this is optimal for visualising gap junction proteins ^63^.

### Immunohistochemistry

Immunohistochemistry was performed as described previously ^19^. Slides with tissue sections were thawed in a benchtop incubator at 37 °C for 30 min to aid the adherence of the tissue to the glass slides and reduce tissue loss during subsequent wash steps. Slides were washed three times in PBS. Hydrophobic barrier pens were used to draw rings around each spinal cord tissue on the slides, and the tissue was then incubated in 1X PBS (pH 7.45, Ca^2+^ and Mg^+^ free, ThermoFisher Scientific, #18912-014) containing 3% Bovine Serum Albumin (BSA; Sigma Aldrich, #A9647) and 0.2% Triton X-100 (Fisher Scientific, #BP151-500) for 2 h at room temperature to block non-specific binding and permeabilise the tissue. Primary antibodies were diluted in 1X PBS containing 1.5% BSA and 0.1% Triton X-100, and samples were incubated with primary antibody solution for two nights at 4 °C. Slides were then washed in 1X PBS five times over the course of 1 h. Secondary antibodies were then added to slides, diluted 1:500 in 0.1% Triton X-100 and incubated for 1.5 h. Slides were washed an additional five times in 1X PBS over the course of an hour. If nuclear labelling was performed, DAPI stain was diluted in PBS at 1:4000 and applied to the slides for 10 min before being washed in DI water to stop the reaction. Finally, the slides were dried, and #1.5 coverslips (VWR, #631-0149) were mounted with Prolong Glass Antifade Mountant (Invitrogen, P36980).

**Table 1.** Antibodies and Nanobodies Used.

| Primary Antibody Target | Species | Dilution | Source | Secondary Antibody Species and Fluorophore |
| --- | --- | --- | --- | --- |
| PSD95-Nb-Atto488 | Camelid | 1:500 | Synaptic Systems, #N3702-At488-L | Atto488 (Conjugated) |
| PSD95-Nb-AlexaFluor647 | Camelid | 1:250 | Synaptic Systems, #N3702-AF647-L | Alexa Fluor 647 (Conjugated) |
| CHAT | Goat | 1:100 | Merck, #AB144P | Donkey Anti-Goat Alexa Fluor 555, Abcam, #ab150134 |
| MMP9 | Goat | 1:500 | Sigma-Aldrich/Merck, #M9570 | Donkey Anti-Goat Alexa Fluor 555, Abcam, #ab150134 |
| VGLUT1-STAR-Orange | Mouse monoclonal | 1:500 | Synaptic Systems, #135 011 | STAR Orange (Conjugated) |
| VGLUT1 | Guinea Pig | 1:500 | Merck, #ab5905 | Donkey Anti-Guinea Pig Alexa Fluor 647, Jackson, #706-605-148 |
| VGLUT2 | Mouse monoclonal | 1:500 | Abcam, #ab79157 | Donkey Anti-Mouse Alexa Fluor 555, Invitrogen, #A31570 |
| Bassoon | Rabbit | 1:500 | Synaptic Systems, #141 003 | Donkey Anti-Rabbit Alexa Fluor 555, Abcam, #ab150062 |
| Cx36 | Mouse monoclonal | 1:500 | ThermoFisher Scientific, #37-4600 | Donkey Anti-Mouse Alexa Fluor 555, Invitrogen, #A31570 |
| ZO-1 | Rabbit Polyclonal | 1:500 | ProteinTech, #21773-1-AP | Donkey Anti-Rabbit Alexa Fluor 555, Abcam, #ab150062 |
| NMDAR2A | Rabbit Polyclonal | 1:500 | ProteinTech, #28571-1-AP | Donkey Anti-Rabbit Alexa Fluor 647 Plus, Invitrogen, #A32795 |
| GluA4 | Rabbit Monoclonal | 1:500 | Cell Signalling, #8070T | Donkey Anti-Rabbit Alexa Fluor 555, Abcam, #ab150062 |
| GluA1 | Rabbit | 1:500 | Abcam, #ab12108 | Donkey Anti-Rabbit Alexa Fluor 555, Abcam, #ab150062 |
| Shank2 | Guinea Pig | 1:250 | Synaptic Systems, #162 204 | Donkey Anti-Guinea Pig Alexa Fluor 647, Jackson, #706-605-148 |

### High-Resolution Confocal Microscopy

High-Resolution microscopy was provided through the Andor BC43 benchtop spinning disk confocal microscope equipped with 405, 488, 561 and 638 nm lasers for illumination. Z-stack images were acquired at a 60 × 1.4 NA oil objective lens, with ×2 frame averaging and a step size of (0.1µm) in accordance with Nyquist criteria. Each channel was acquired sequentially in the Z-plane.

### Airyscan Microscopy

High-resolution Airyscan microscopy was performed using the Zeiss LSM800 laser scanning confocal microscope, based on an Axio ‘Observer 7’ microscope, equipped with a 63 × 1.4 NA oil objective lens, an Airyscan super-resolution module plus two individual GaAsP PMT detectors. Illumination was provided by 405, 488, 561 and 640 nm laser lines. The pixel size was set to 0.04 μm to provide optimal resolution. A minimum of 2× line averaging was performed in unidirectional scanning acquisitions. Z-stacks were acquired with 150 nm step size in accordance with Nyquist sampling rate, with each channel acquired sequentially in the Z-plane. Airyscan processing on images was performed in 3D to yield sub-diffraction-limit images of structures, with the Wiener filter kept consistent for images within a data set (value of 6 for VGLUT1, value of 7 for PSD95 and other synaptic markers). Images were acquired in the LMC of the ventral horn.

### SIM^2^ Microscopy

Z-stack images were acquired using a Zeiss Elyra 7 LatticeSIM microsystem with a Plan-Apochromat 63 × 1.4 NA oil objective using the 488 nm (at 5.5%) and 561 nm (at 5.5%) 500 mW lasers in LatticeSIM mode, capturing 13 phases with the 32.00 µm Grating selected. Images (Dimensions: x: 2560, y: 2560, z: 147, channels: 2, 16-bit) underwent SIM^2^ post-acquisition processing, SIM^2^ settings ‘weak fixed’ with parameters adjusted for 5 iterations and 0 regularisation.

### g-STED Microscopy

Multi-channel confocal and gated-stimulated emission depletion (g-STED) microscopy was performed using the Leica SP8 SMD g-STED microscope available at the Edinburgh Super-Resolution Imaging Consortium (ESRIC) hosted by Heriot Watt University. Excitation was provided by a CW super-continuum white light laser source (12 – 30 % of maximum power). Depletion was provided by a 594 nm laser using the 3D Pi depletion (2D STED laser: 17 – 25% of maximum power; 3D STED laser: 12 – 30% of maximum power). Images were acquired with a 100 × 1.4 NA oil STED objective lens. The optical zoom was set to achieve a resultant image pixel size of 20 nm. ×3 line averaging was performed in unidirectional scanning acquisitions at a scan speed of 700 – 1000 Hz. Fluorescence was detected using a Leica Hybrid detector, minimum time gate at 1 or 1.2 ns, and a maximum time gating of 8 ns. Confocal and g-STED images were captured sequentially (confocal then g-STED).

### Imaris Analysis

IMARIS (Imaris Version 9.9.1) was used to detect and quantify the number and structure of Honeycomb Synapses and their spatial colocalisation with VGLUT1 and MN markers from the high-resolution confocal images. 3D reconstructions (Surfaces) of immunohistochemically or genetically labelled presynaptic markers (Pv-Cre:YFP, VGLUT1), postsynaptic marker PSD95 and MN markers (MMP9 or CHAT) were generated from z-stack images to enable quantitative analysis of their size and colocalization with different markers. The Surface creation tool was used to render each marker.

Smoothing and background subtraction (local contrast) were applied, with parameters chosen so that the resulting 3D outline best reflects the raw data. 3D outlines were then segmented into individual objects using automatic thresholding. A minimum voxel count filter was applied to omit small objects, reducing background noise. Individual MNs were manually identified from MMP9 objects based on their distinct morphology and labelled using the ‘object classification’ option in Imaris. The volume of each object was measured and the shortest distance of PSD95-labelled objects to the nearest presynaptic and MN marker was calculated.

A supervised, built-in Machine Learning Algorithm (MLA) was trained in IMARIS to identify postsynaptic Honeycomb (HC-PSD) and non-HC (NHC-PSD) PSDs. The MLA was trained using manually labelled examples of HC- and NHC-PSDs from 26 images. Subsequently, the MLA was able to assign each PSD95 object to the ‘HC-PSD’ or ‘NHC-PSD’ class, based on a multivariate feature space including morphological parameters. All predictions from this MLA were manually checked and corrected if appropriate, with the MLA acting only as a tool to improve the accuracy and efficiency of of Honeycomb Synapse detection.

Researchers analysing data to investigate synapse pathology in ALS models were blinded to the genotype and disease state of the slides during imaging and analysis. Blinding was not used for analysis of Honeycomb Synapse nanostructure or expression across different anatomical regions or development.

### FIJI Image Analysis

Image analysis of Honeycomb Synapse nanostructure and molecular components was conducted in FIJI ^92^. Z-stack acquisitions were converted to 2D through an average projection to capture the PSD structure. SIM^2^ images required no pre-processing. g-STED images underwent background subtraction (14-pixel radius) and Gaussian smoothing (2-pixel radius). The PSD matrix was detected by manual thresholding. The ‘fill holes’ function was used to discern the area of the PSD including and excluding the perforations. The number of complete perforations was determined manually using the Count tool. The number of PSD95 clusters around each perforation was determined semi-automatedly by using the Find Maxima function to best detect each intensity peak surrounding each perforation (SIM^2^ images using a maxima prominence of 40 – 100 across images; g-STED images using a maxima prominence of 5 – 15 across the images).

For the analysis of Honeycomb Synapse molecular components from Airyscan image acquisitions, images PSD95 and other synaptic markers were processed using a background subtraction (12-pixel radius). Each marker was manually thresholded. For differential detection of PSD95 subdomains, the matrix of the PSD was detected by manually thresholding the PSD95 signal. To detect the perforations, the whole PSD was thresholded including the perforations (using the Fill Holes function), and the Image Calculator tool was used to produce the difference between the whole PSD and the PSD matrix images – providing only the PSD perforations binarized. JACoP (Just Another Colocalisation Plugin ^93^) was used to measure the spatial colocalization (Mander’s Coefficient) of synaptic markers with both the matrix and the perforations of the Honeycomb Synapses.

### Statistics

For the analysis of molecular components and their colocalization within the PSD95 Honeycomb Synapse domains (Figure 3), a one-way ANOVA followed by Tukey’s HSD post-hoc test was performed in Graphpad Prism (version 10.6.1) to identify differences between synaptic markers in their association with the Honeycomb Synapse matrix and perforations. In the developmental data set (Figure 4), a one-way ANOVA followed by Tukey’s HSD post-hoc test was performed to identify differences in postnatal age groups. In the anatomical mapping data (Figure 5), analysis was performed using Python (version 3.13.3) and stored in Jupyter Notebooks (version 2025.7.0).

Statistical analyses were conducted with the packages statsmodels (version 0.14.4), scipy.stats (version 1.15.2) and patsy (version 1.0.1). Anatomical mapping data were analysed using generalised linear models (GLMs) and linear mixed-effect models (LMMs). To account for repeated measures from the same animal (n=3) all models used clustered robust standard errors or random intercepts for each animal. Logistic regression models assuming a binomial distribution were fitted to analyse binary outcomes. Poisson regression models were fitted to model count data. If overdispersion was detected in count data, negative binomial regression models were used. LMMs were fitted to analyse continuous dependent variables. For the analysis of Honeycomb Synapse structure in plasticity experiments (Figure 7), a one-way Welch’s ANOVA with Games-Howell post-hoc test was performed to determine differences in Honeycomb Synapse PSD volume between sham, anodal, and cathodal stimulation. For the analysis of Honeycomb Synapses in ALS models compared with controls (Figure 8), two-sample T-tests were performed in Graphpad Prism. Shapiro-Wilk tests were performed to assess normality, and a Levene’s test was used to assess the variance equality of distribution (SPSS or Graphpad Prism). Data was collated and charts prepared either using Microsoft Excel or Graphpad Prism, typically denoting the mean ± standard error of the mean along with individual data points. Unless otherwise stated, statistical significance in graphs is denoted as follows: * p<0.05, ** p<0.01, *** p<0.001, **** p<0.0001.

## Supplementary Figures

**Supplementary Figure 1.**
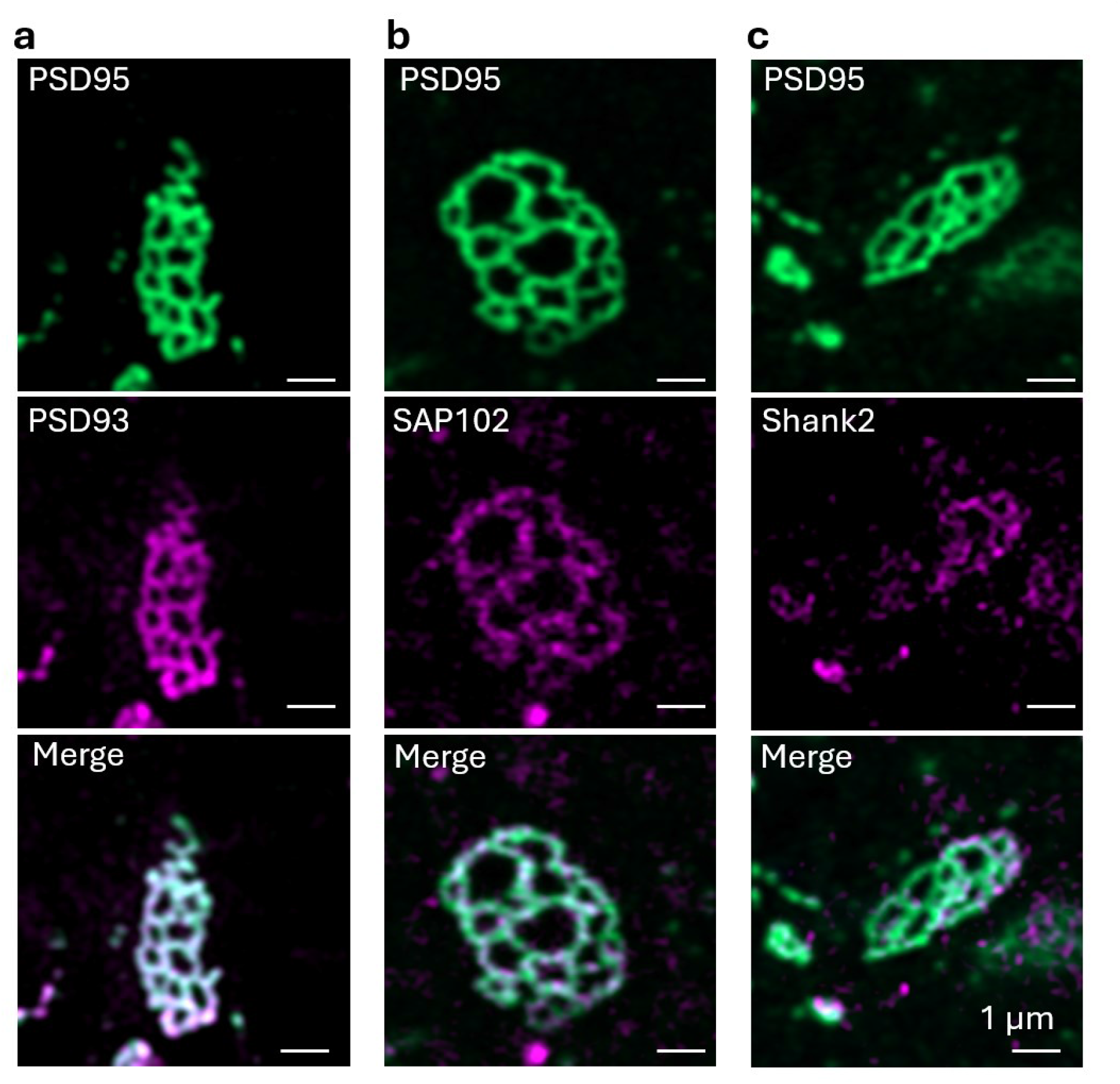
Organisation of Scaffold Proteins in Honeycomb Synapses. PSD95-eGFP Honeycomb Synapses were visualised, alongside PSD93 (a), SAP102 (b) and Shank2 (c), showing the separate images, and the merge composite.

**Supplementary Figure 2.**
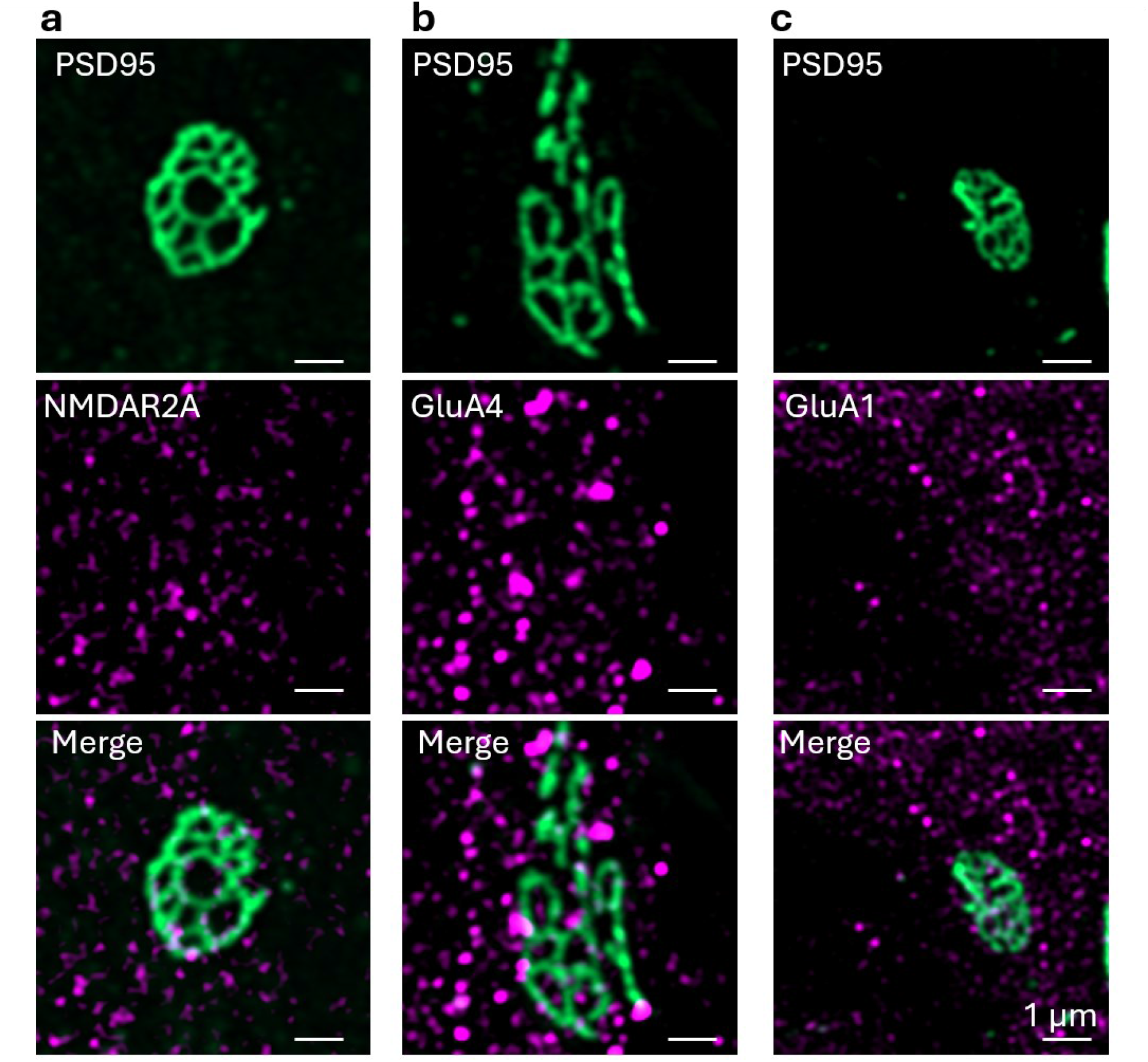
Organisation of Glutamatergic Receptors in Honeycomb Synapses. PSD95-eGFP Honeycomb Synapses were visualised, alongside NMDAR2A (a), GluA4 (b) and GluA1 (c), showing the separate images, and the merge composite.

**Supplementary Figure 3.**
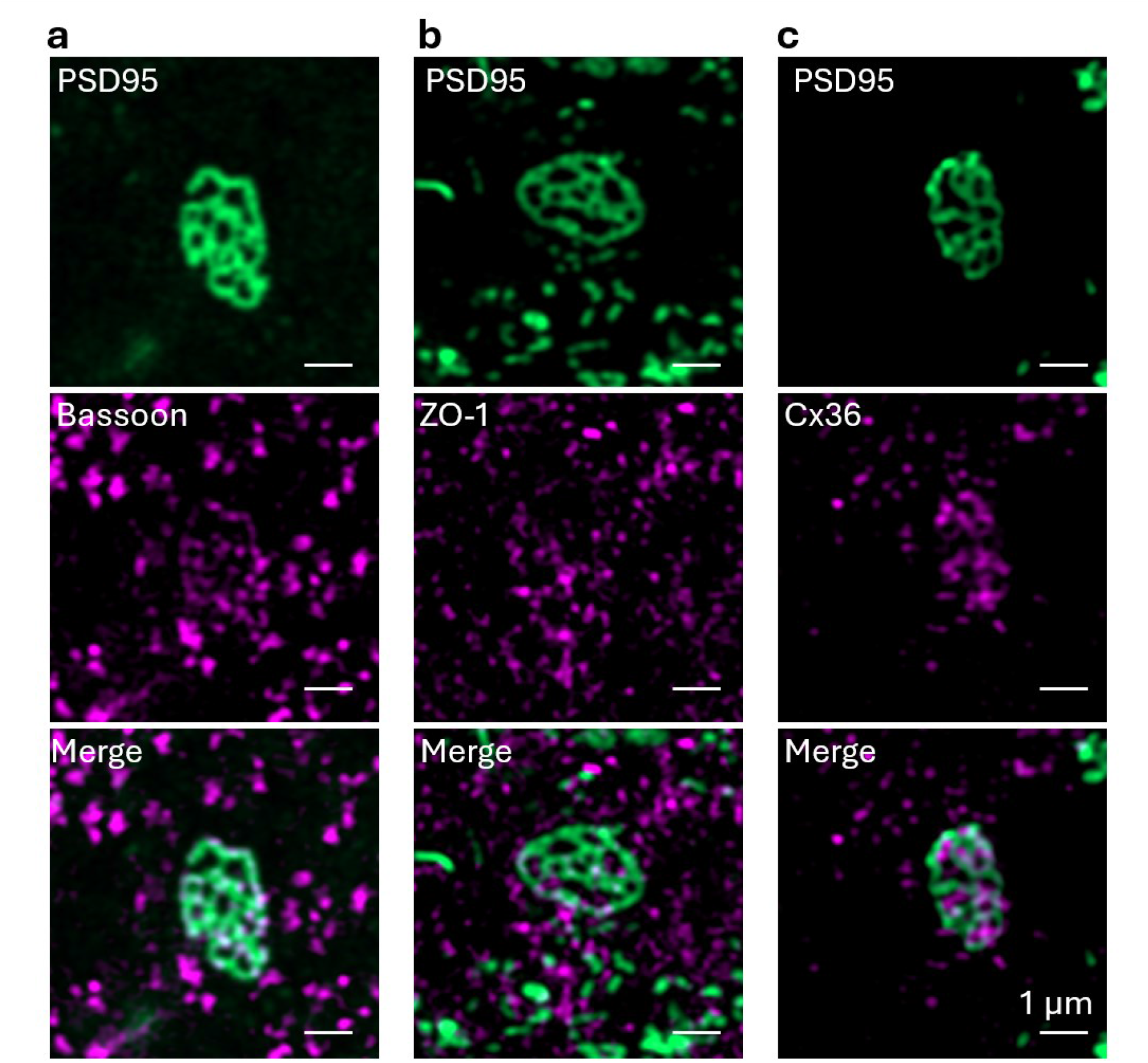
Organisation of Vesicular Release and Gap Junction Proteins in Honeycomb Synapses. PSD95-eGFP Honeycomb Synapses were visualised, alongside Bassoon (a), ZO-1 (b) and Cx36 (c), showing the separate images, and the merge composite.

**Supplementary Figure 4.**
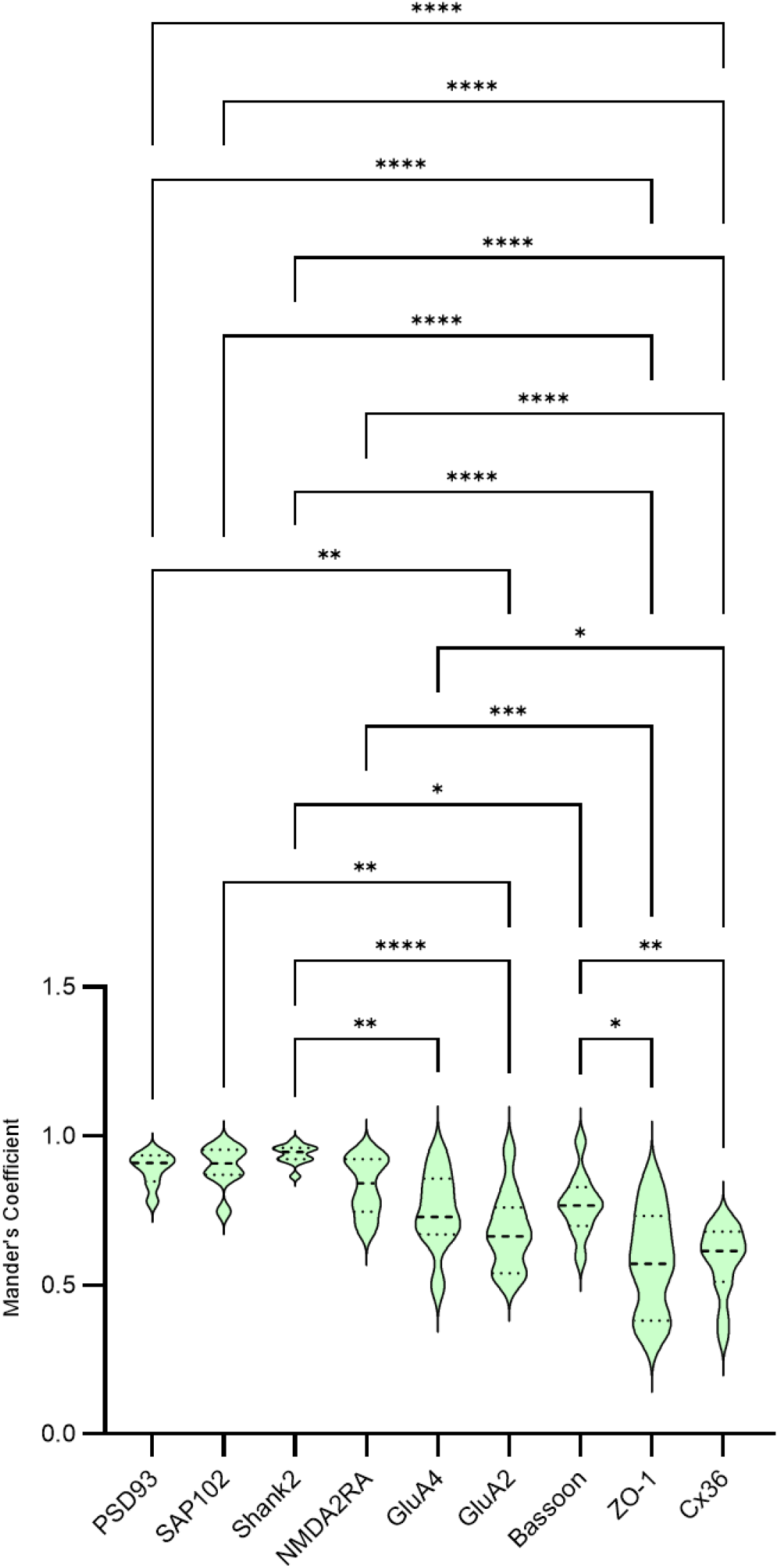
Bar chart plotting the degree of colocalization (Mander’s Coefficient) of each synaptic marker within the Honeycomb Synapse Matrix as discerned from PSD95-eGFP expression. * = p<0.05, **p<0.01, *** p<0.001, **** p<0.0001

**Supplementary Figure 5.**
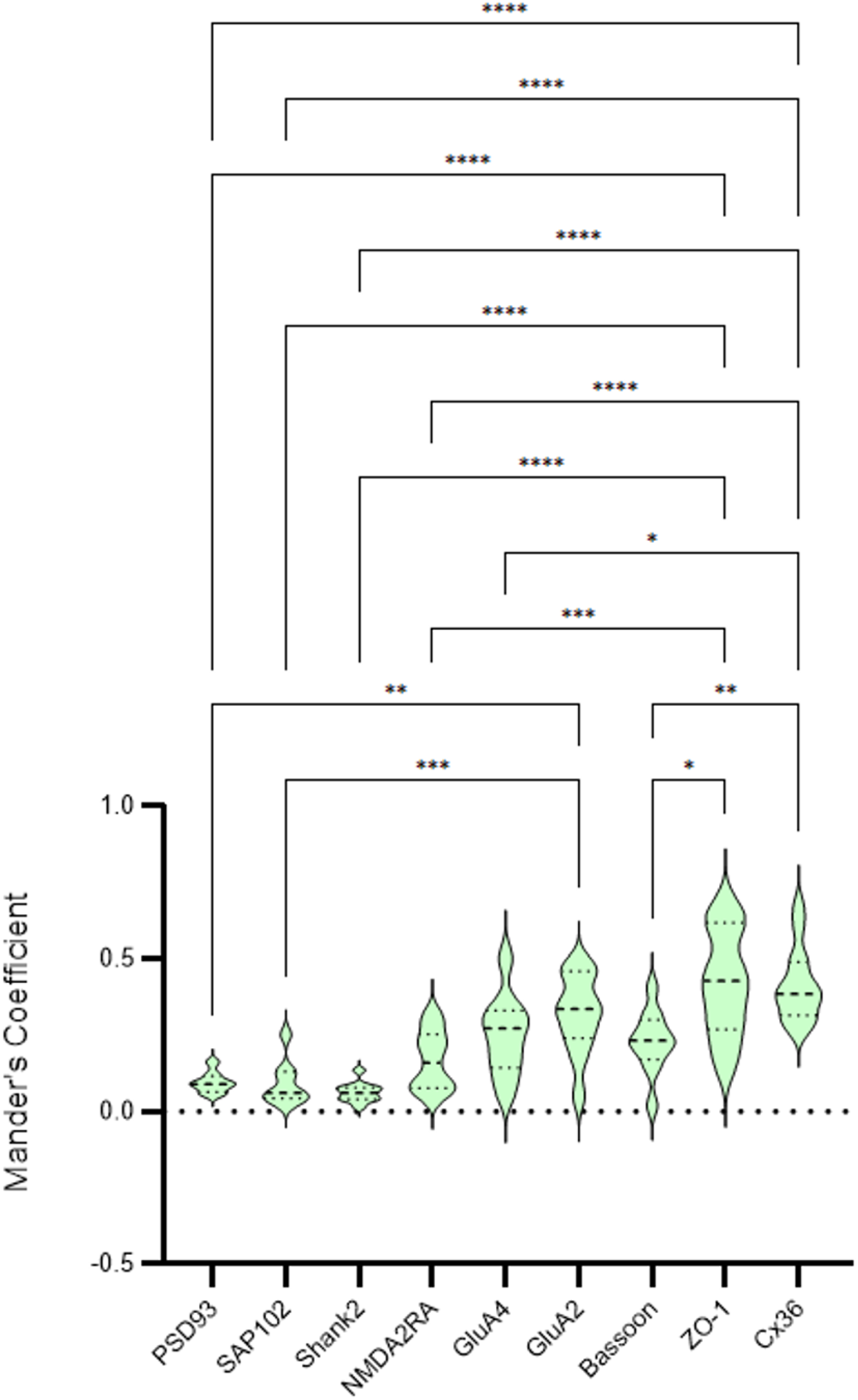
Bar chart plotting the degree of colocalization (Mander’s Coefficient) of each synaptic marker within the Honeycomb Synapse perforations as discerned from PSD95-eGFP expression. * = p<0.05, **p<0.01, ***p<0.001, ****p<0.0001

**Supplementary Figure 6.**
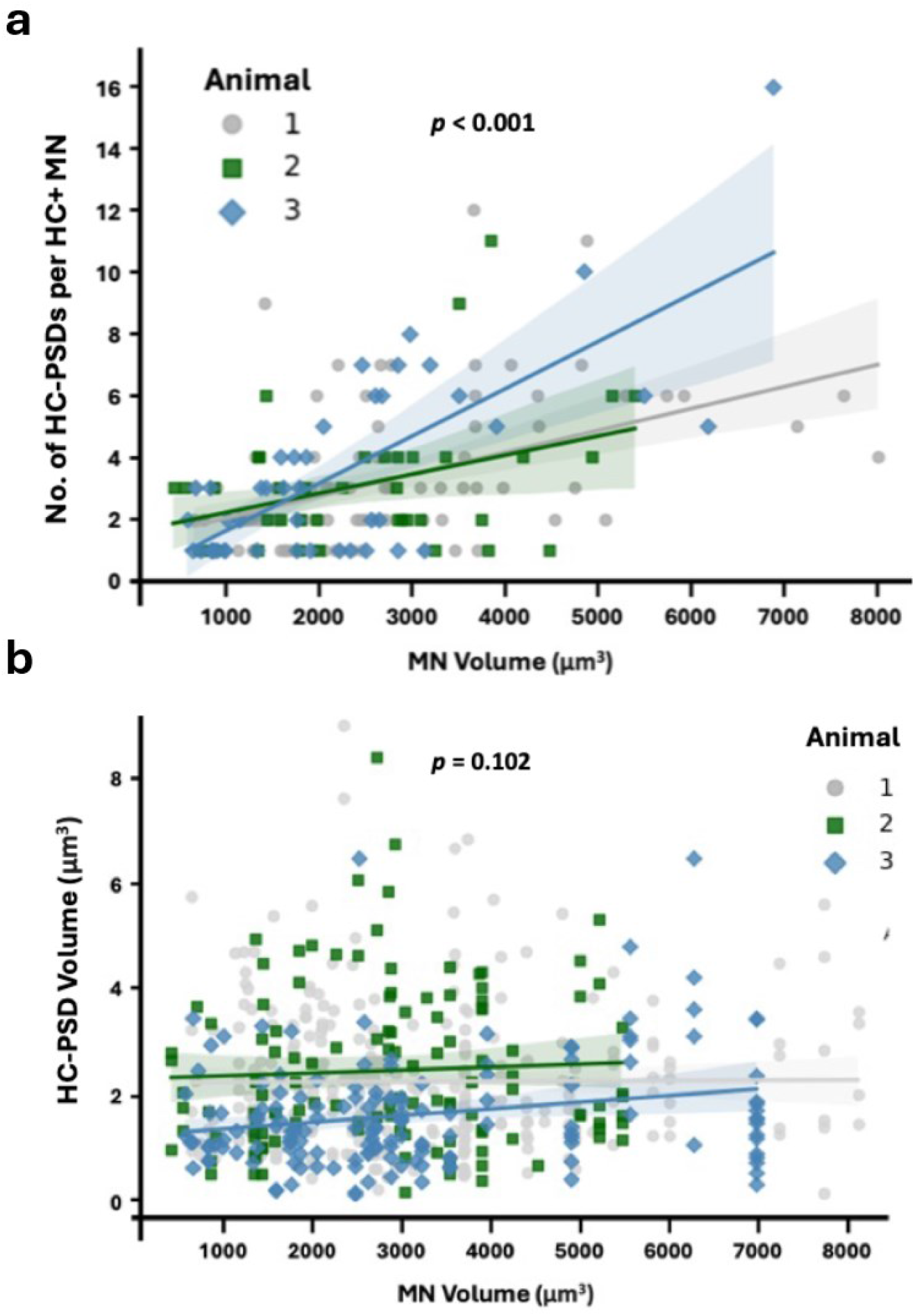
a, Scatter plot showing number of colocalised Honeycomb Synapse postsynaptic densities (HC-PSD) per the volume of the motoneuron (MN) that harboured Honeycomb Synapses (HC+ MN). The data demonstrates a positive correlation between MN size and the number of Honeycomb Synapses harboured by the MN. b, Scatter plot showing HC-PSD volume as a function of MN volume by animal. The data demonstrates there is no correlation between MN size and Honeycomb Synapse size. Data was separated by each of the 3 animals as per the keys to the right of each graph.

**Supplementary Figure 7.**
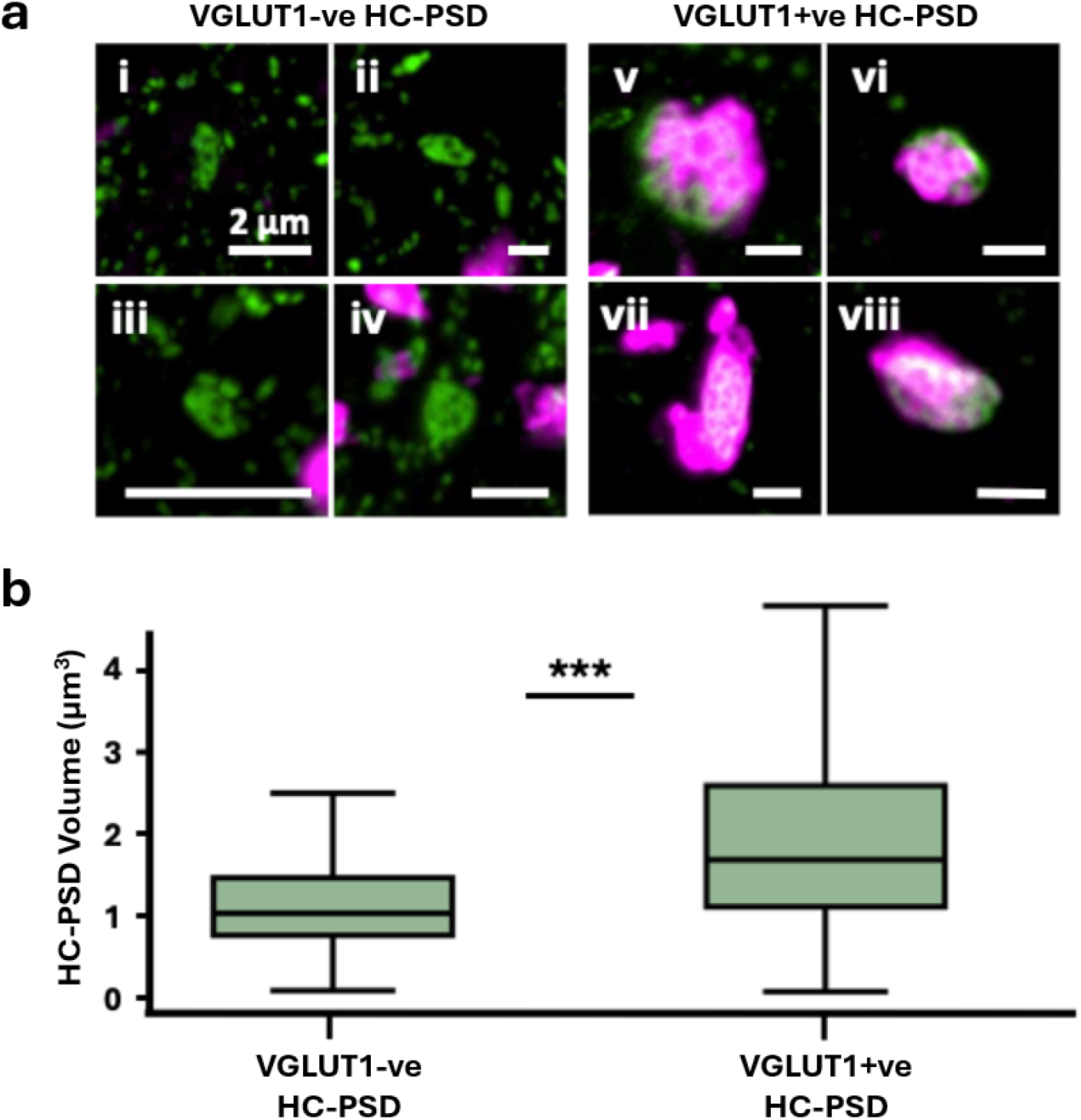
a, Honeycomb Synapses were identified from PSD95 structures harbouring at least 2 perforations. Whilst most Honeycomb Synapses were observed opposed to VGLUT1 (VGLUT1+ve HC-PSD; examples i-iv), a small portion were not (VGLUT1-ve HC-PSD; examples v-viii). b, Graph plotting the size of VGLUT1 -ve and VGLUT1 +ve HC-PSDs. VGLUT1 +ve HC-PSDs are significantly larger than VGLUT1 -ve HC- PSDs ((3 = 0.746, p < 0.001).

## References.

1. Grant, S. G. N. & Fransén, E. The Synapse Diversity Dilemma: Molecular Heterogeneity Confounds Studies of Synapse Function. Front Synaptic Neurosci 12, 590403 (2020).

2. Emes, R. D. & Grant, S. G. N. Evolution of synapse complexity and diversity. Annu Rev Neurosci 35, 111–131 (2012).

3. Emes, R. D., et al. Evolutionary expansion and anatomical specialization of synapse proteome complexity. Nat Neurosci 11, 799–806 (2008).

4. Connor, S. A. & Siddiqui, T. J. Synapse organizers as molecular codes for synaptic plasticity. Trends Neurosci 46, 971–985 (2023).

5. Südhof, T. C. & Malenka, R. C. Understanding Synapses: Past, Present, and Future. Neuron 60, 469–476 (2008).

6. Tang, A.-H., et al. A trans-synaptic nanocolumn aligns neurotransmitter release to receptors. Nature 536, 210–214 (2016).

7. Frank, R. A. & Grant, S. G. Supramolecular organization of NMDA receptors and the postsynaptic density. Current Opinion in Neurobiology 45, 139–147 (2017).

8. Benavides-Piccione, R., Fernaud-Espinosa, I., Kastanauskaite, A. & DeFelipe, J. Principles for Dendritic Spine Size and Density in Human and Mouse Cortical Pyramidal Neurons. J Comp Neurol 533, e70060 (2025).

9. Schünemann, K. D., et al. Comprehensive analysis of human dendritic spine morphology and density. J Neurophysiol 133, 1086–1102 (2025).

10. MacGillavry, H. D., Song, Y., Raghavachari, S. & Blanpied, T. A. Nanoscale scaffolding domains within the postsynaptic density concentrate synaptic AMPA receptors. Neuron 78, 615–622 (2013).

11. Nair, D., et al. Super-resolution imaging reveals that AMPA receptors inside synapses are dynamically organized in nanodomains regulated by PSD95. J Neurosci 33, 13204–13224 (2013).

12. Broadhead, M. J., et al. PSD95 nanoclusters are postsynaptic building blocks in hippocampus circuits. Sci Rep 6, 24626 (2016).

13. Jafri, H., Thomas, S. J., Yang, S. H., Cain, R. E. & Dalva, M. B. Nano-organization of synapses defines synaptic release properties at cortical neuron dendritic spines. bioRxiv 2025.02.13.637710 (2025) doi:10.1101/2025.02.13.637710.

14. Sherrington, C. S. The Integrative Action of the Nervous System. in Scientific and Medical Knowledge Production, 1796-1918 (Routledge, 2023).

15. Bernstein, J. J. & Bernstein, M. E. Ventral horn synaptology in the rat. J Neurocytol 5, 109–123 (1976).

16. Conradi, S. Functional Implications of Structure and Synaptology of Motor Neurons in Motor Neuron Disease. in Clinical Aspects of Sensory Motor Integration (eds Struppler, A. & Weindl, A.) 86–90 (Springer, Berlin, Heidelberg, 1987). doi:10.1007/978-3-642-71540-2_10.

17. Tahayori, B. & Koceja, D. M. Activity-Dependent Plasticity of Spinal Circuits in the Developing and Mature Spinal Cord. Neural Plast 2012, 964843 (2012).

18. Wishart, T. M., Parson, S. H. & Gillingwater, T. H. Synaptic vulnerability in neurodegenerative disease. J Neuropathol Exp Neurol 65, 733–739 (2006).

19. Ayvazian-Hancock, A., Butler, E., Meehan, C. F., Miles, G. B. & Broadhead, M. J. Synaptopathy in the TDP-43ΔNLS Mouse Model of Sporadic Amyotrophic Lateral Sclerosis. Eur J Neurosci 62, e70320 (2025).

20. Broadhead, M. J., et al. Selective vulnerability of tripartite synapses in amyotrophic lateral sclerosis. Acta Neuropathol 143, 471–486 (2022).

21. Sangari, S., Lackmy-Vallée, A., Peyre, I., Pradat, P.-F. & Marchand-Pauvert, V. Afferent-driven modulation of spinal interneuron circuits across disease stages in amyotrophic lateral sclerosis. Neurobiology of Disease 219, 107270 (2026).

22. Fogarty, M. J. Amyotrophic lateral sclerosis as a synaptopathy. Neural Regen Res 14, 189–192 (2019).

23. Devlin, A.-C., et al. Human iPSC-derived motoneurons harbouring TARDBP or C9ORF72 ALS mutations are dysfunctional despite maintaining viability. Nat Commun 6, 5999 (2015).

24. Suzuki, K., et al. A synthetic synaptic organizer protein restores glutamatergic neuronal circuits. Science 369, eabb4853 (2020).

25. Mora, S., et al. Stabilization of V1 interneuron-motor neuron connectivity ameliorates motor phenotype in a mouse model of ALS. Nat Commun 15, 4867 (2024).

26. Zhu, F., et al. Architecture of the Mouse Brain Synaptome. Neuron 99, 781–799.e10 (2018).

27. Broadhead, M. J., et al. Nanostructural Diversity of Synapses in the Mammalian Spinal Cord. Sci Rep 10, 8189 (2020).

28. Leitch, B., Cobb, J. L., Heitler, W. J. & Pitman, R. M. Post-embryonic development of rectifying electrical synapses in the crayfish: ultrastructure. J Neurocytol 18, 749–761 (1989).

29. Brown, A. G. & Fyffe, R. E. Direct observations on the contacts made between Ia afferent fibres and alpha-motoneurones in the cat’s lumbosacral spinal cord. J Physiol 313, 121–140 (1981).

30. Alvarez, F. J., Villalba, R. M., Zerda, R. & Schneider, S. P. Vesicular glutamate transporters in the spinal cord, with special reference to sensory primary afferent synapses. J Comp Neurol 472, 257–280 (2004).

31. Jankowiak, T., Cholewiński, M. & Bączyk, M. Differential Effects of Invasive Anodal Trans-spinal Direct Current Stimulation on Monosynaptic Excitatory Postsynaptic Potentials, Ia Afferents Excitability, and Motoneuron Intrinsic Properties Between Superoxide Dismutase Type-1 Glycine to Alanine Substitution at Position 93 and Wildtype Mice. Neuroscience 498, 125–143 (2022).

32. Andersen, P. M. & Al-Chalabi, A. Clinical genetics of amyotrophic lateral sclerosis: what do we really know? Nat Rev Neurol 7, 603–615 (2011).

33. Bączyk, M., et al. Synaptic restoration by cAMP/PKA drives activity-dependent neuroprotection to motoneurons in ALS. J Exp Med 217, e20191734 (2020).

34. Allodi, I., Montañana-Rosell, R., Selvan, R., Löw, P. & Kiehn, O. Locomotor deficits in a mouse model of ALS are paralleled by loss of V1-interneuron connections onto fast motor neurons. Nat Commun 12, 3251 (2021).

35. Nagao, M., Misawa, H., Kato, S. & Hirai, S. Loss of cholinergic synapses on the spinal motor neurons of amyotrophic lateral sclerosis. J Neuropathol Exp Neurol 57, 329–333 (1998).

36. Alvarez, F. J., Bullinger, K. L., Titus, H. E., Nardelli, P. & Cope, T. C. Permanent reorganization of Ia afferent synapses on motoneurons after peripheral nerve injuries. Ann N Y Acad Sci 1198, 231– 241 (2010).

37. Broadhead, M. J., et al. Synaptic expression of TAR-DNA-binding protein 43 in the mouse spinal cord determined using super-resolution microscopy. Front Mol Neurosci 16, 1027898 (2023).

38. Nascimento, F., et al. Spinal microcircuits go through multiphasic homeostatic compensations in a mouse model of motoneuron degeneration. Cell Rep 43, 115046 (2024).

39. Mancuso, R., Santos-Nogueira, E., Osta, R. & Navarro, X. Electrophysiological analysis of a murine model of motoneuron disease. Clinical Neurophysiology 122, 1660–1670 (2011).

40. Gurney, M. E. Transgenic-mouse model of amyotrophic lateral sclerosis. N Engl J Med 331, 1721– 1722 (1994).

41. Vinsant, S., et al. Characterization of early pathogenesis in the SOD1G93A mouse model of ALS: part II, results and discussion. Brain Behav 3, 431–457 (2013).

42. Walker, A. K., et al. Functional recovery in new mouse models of ALS/FTLD after clearance of pathological cytoplasmic TDP-43. Acta Neuropathol 130, 643–660 (2015).

43. Bak, A. N., et al. Cytoplasmic TDP-43 accumulation drives changes in C-bouton number and size in a mouse model of sporadic Amyotrophic Lateral Sclerosis. Mol Cell Neurosci 125, 103840 (2023).

44. Fukata, Y., et al. Local palmitoylation cycles define activity-regulated postsynaptic subdomains. J Cell Biol 202, 145–161 (2013).

45. Metzbower, S. R., Levy, A. D., Dharmasri, P. A., Anderson, M. C. & Blanpied, T. A. Distinct SAP102 and PSD-95 Nano-organization Defines Multiple Types of Synaptic Scaffold Protein Domains at Single Synapses. J Neurosci 44, e1715232024 (2024).

46. Gonzales, R. B., DeLeon Galvan, C. J., Rangel, Y. M. & Claiborne, B. J. Distribution of thorny excrescences on CA3 pyramidal neurons in the rat hippocampus. J Comp Neurol 430, 357–368 (2001).

47. Rodríguez-Contreras, A., van Hoeve, J. S. S., Habets, R. L. P., Locher, H. & Borst, J. G. G. Dynamic development of the calyx of Held synapse. Proceedings of the National Academy of Sciences 105, 5603–5608 (2008).

48. Paradiso, K., Wu, W. & Wu, L.-G. Methods for patch clamp capacitance recordings from the calyx. J Vis Exp 244 (2007) doi:10.3791/244.

49. Wang, L.-Y., Neher, E. & Taschenberger, H. Synaptic Vesicles in Mature Calyx of Held Synapses Sense Higher Nanodomain Calcium Concentrations during Action Potential-Evoked Glutamate Release. J. Neurosci. 28, 14450–14458 (2008).

50. Baydyuk, M., Xu, J. & Wu, L.-G. The Calyx of Held in the auditory system: structure, function, and development. Hear Res 338, 22–31 (2016).

51. Reschechtko, S. & Pruszynski, J. A. Stretch reflexes. Current Biology 30, R1025–R1030 (2020).

52. Theodosiadou, A., Henry, M., Duchateau, J. & Baudry, S. Revisiting the use of Hoffmann reflex in motor control research on humans. Eur J Appl Physiol 123, 695–710 (2023).

53. Magladery, J. W. & McDOUGAL, D. B. Electrophysiological studies of nerve and reflex activity in normal man. I. Identification of certain reflexes in the electromyogram and the conduction velocity of peripheral nerve fibers. Bull Johns Hopkins Hosp 86, 265–290 (1950).

54. Lazar, J. W. The early history of the knee-jerk reflex in neurology. J Hist Neurosci 31, 409–424 (2022).

55. Pierce, J. P. & Mendell, L. M. Quantitative ultrastructure of Ia boutons in the ventral horn: scaling and positional relationships. J Neurosci 13, 4748–4763 (1993).

56. Harris, K. M. & Weinberg, R. J. Ultrastructure of synapses in the mammalian brain. Cold Spring Harb Perspect Biol 4, a005587 (2012).

57. Lovatt, C., O’Sullivan, T., Ortega-de San Luis, C., Ryan, T. J. & Frank, R. A. W. Memory engram synapse 3D macromolecular architecture visualized by cryoCLEM-guided cryoET. Structure 34, 100–112.e3 (2026).

58. Yang, X. & Specht, C. G. Subsynaptic Domains in Super-Resolution Microscopy: The Treachery of Images. Front Mol Neurosci 12, 161 (2019).

59. Nieto-Sampedro, M., Hoff, S. F. & Cotman, C. W. Perforated postsynaptic densities: probable intermediates in synapse turnover. Proc Natl Acad Sci U S A 79, 5718–5722 (1982).

60. Bautista, W., Nagy, J. I., Dai, Y. & McCrea, D. A. Requirement of neuronal connexin36 in pathways mediating presynaptic inhibition of primary afferents in functionally mature mouse spinal cord. J Physiol 590, 3821–3839 (2012).

61. Nagy, J. I., Bautista, W., Blakley, B. & Rash, J. E. Morphologically mixed chemical-electrical synapses formed by primary afferents in rodent vestibular nuclei as revealed by immunofluorescence detection of connexin36 and vesicular glutamate transporter-1. Neuroscience 252, 468–488 (2013).

62. Nagy, J. I., Lynn, B. D., Senecal, J. M. M. & Stecina, K. Connexin36 Expression in Primary Afferent Neurons in Relation to the Axon Reflex and Modality Coding of Somatic Sensation. Neuroscience 383, 216–234 (2018).

63. Bautista, W., McCrea, D. A. & Nagy, J. I. Connexin36 identified at morphologically mixed chemical/electrical synapses on trigeminal motoneurons and at primary afferent terminals on spinal cord neurons in adult mouse and rat. Neuroscience 263, 159–180 (2014).

64. Silwal, P., et al. Patterns of connexin36 and eGFP reporter expression among motoneurons in spinal sexually dimorphic motor nuclei in mouse. Int J Physiol Pathophysiol Pharmacol 16, 55–76 (2024).

65. Moreno, A. P. & Lau, A. F. Gap junction channel gating modulated through protein phosphorylation. Prog Biophys Mol Biol 94, 107–119 (2007).

66. Pereda, A. E. & Miller, A. C. Defining the electrical synapse. Nat Rev Neurosci 10.1038/s41583-026-01080-y (2026).

67. Bhattacharya, A., Aghayeva, U., Berghoff, E. G. & Hobert, O. Plasticity of the Electrical Connectome of C. elegans. Cell 176, 1174–1189.e16 (2019).

68. Peracchia, C. Calcium Role in Gap Junction Channel Gating: Direct Electrostatic or Calmodulin-Mediated? Int J Mol Sci 25, 9789 (2024).

69. Kita, K., et al. GluA4 facilitates cerebellar expansion coding and enables associative memory formation. Elife 10, e65152 (2021).

70. Vega-Gutiérrez, C., et al. GluA4 AMPA receptor gating mechanisms and modulation by auxiliary proteins. Nat Struct Mol Biol 1–13 (2025).

71. Erreger, K., Dravid, S. M., Banke, T. G., Wyllie, D. J. A. & Traynelis, S. F. Subunit-specific gating controls rat NR1/NR2A and NR1/NR2B NMDA channel kinetics and synaptic signalling profiles. The Journal of Physiology 563, 345–358 (2005).

72. Glasgow, N. G., Siegler Retchless, B. & Johnson, J. W. Molecular bases of NMDA receptor subtype-dependent properties. J Physiol 593, 83–95 (2015).

73. Sharples, S. A., Broadhead, M. J., Gray, J. A. & Miles, G. B. M-type potassium currents differentially affect activation of motoneuron subtypes and tune recruitment gain. J Physiol 601, 5751–5775 (2023).

74. McHanwell, S. & Biscoe, T. J. The sizes of motoneurons supplying hindlimb muscles in the mouse. Proc R Soc Lond B Biol Sci 213, 201–216 (1981).

75. Alvarez, F. J. Gephyrin and the regulation of synaptic strength and dynamics at glycinergic inhibitory synapses. Brain Res Bull 129, 50–65 (2017).

76. Bączyk, M., Krutki, P. & Zytnicki, D. Is there hope that transpinal direct current stimulation corrects motoneuron excitability and provides neuroprotection in amyotrophic lateral sclerosis? Physiol Rep 9, e14706 (2021).

77. Wang, G., Gilbert, J. & Man, H.-Y. AMPA Receptor Trafficking in Homeostatic Synaptic Plasticity: Functional Molecules and Signaling Cascades. Neural Plast 2012, 825364 (2012).

78. Mentis, G. Z., et al. Early functional impairment of sensory-motor connectivity in a mouse model of spinal muscular atrophy. Neuron 69, 453–467 (2011).

79. Vaughan, S. K., Kemp, Z., Hatzipetros, T., Vieira, F. & Valdez, G. Degeneration of proprioceptive sensory nerve endings in mice harboring amyotrophic lateral sclerosis-causing mutations. J Comp Neurol 523, 2477–2494 (2015).

80. Montañana-Rosell, R., et al. Spinal inhibitory neurons degenerate before motor neurons and excitatory neurons in a mouse model of ALS. Sci Adv 10, eadk3229 (2024).

81. Scekic-Zahirovic, J., et al. Cytoplasmic FUS triggers early behavioral alterations linked to cortical neuronal hyperactivity and inhibitory synaptic defects. Nat Commun 12, 3028 (2021).

82. Wells, T. L., Myles, J. R. & Akay, T. C-Boutons and Their Influence on Amyotrophic Lateral Sclerosis Disease Progression. J Neurosci 41, 8088–8101 (2021).

83. Herron, L. R. & Miles, G. B. Gender-specific perturbations in modulatory inputs to motoneurons in a mouse model of amyotrophic lateral sclerosis. Neuroscience 226, 313–323 (2012).

84. Wong, C.-E., et al. TDP-43 proteinopathy impairs mRNP granule mediated postsynaptic translation and mRNA metabolism. Theranostics 11, 330–345 (2021).

85. Jensen, D. B., Kadlecova, M., Allodi, I. & Meehan, C. F. Spinal motoneurones are intrinsically more responsive in the adult G93A SOD1 mouse model of amyotrophic lateral sclerosis. J Physiol 598, 4385–4403 (2020).

86. Morozova, V., Hassieb, M. & Ahmed, Z. Multi-site Anodal Direct Current Stimulation Preserves Motor Function and Modulates Key Cellular Pathways in TDP-43 and SOD1 Mouse Models of ALS. Neuromodulation: Technology at the Neural Interface https://doi.org/10.1016/j.neurom.2026.08.007 (2026) doi:10.1016/j.neurom.2026.08.007.

87. Bulovaite, E., et al. A brain atlas of synapse protein lifetime across the mouse lifespan. Neuron 110, 4057–4073.e8 (2022).

88. Fernández, E., et al. Targeted tandem affinity purification of PSD-95 recovers core postsynaptic complexes and schizophrenia susceptibility proteins. Mol Syst Biol 5, 269 (2009).

89. Jankowiak, T., et al. Increase in Ia Afferent Synaptic Excitation of SOD1 G93A Mouse Motoneurons by 2-Week Anodal Trans-Spinal Direct Current Stimulation Does Not Ameliorate the Cellular Burden of the Disease. Eur J Neurosci 62, e70375 (2025).

90. de Oliveira Pires, L., et al. A computational model of tsDCS effects in SOD1 mice: from MRI-based design to validation. Comput Biol Med 197, 111082 (2025).

91. Fernandes, S. R., Salvador, R., Wenger, C., de Carvalho, M. & Miranda, P. C. Transcutaneous spinal direct current stimulation of the lumbar and sacral spinal cord: a modelling study. J Neural Eng 15, 036008 (2018).

92. Schindelin, J., et al. Fiji: an open-source platform for biological-image analysis. Nat Methods 9, 676–682 (2012).

93. Bolte, S. & Cordelières, F. P. A guided tour into subcellular colocalization analysis in light microscopy. J Microsc 224, 213–232 (2006).

