## Supplementary Figures for "Discovery of the Honeycomb Synapse in Spinal Motor Circuits"

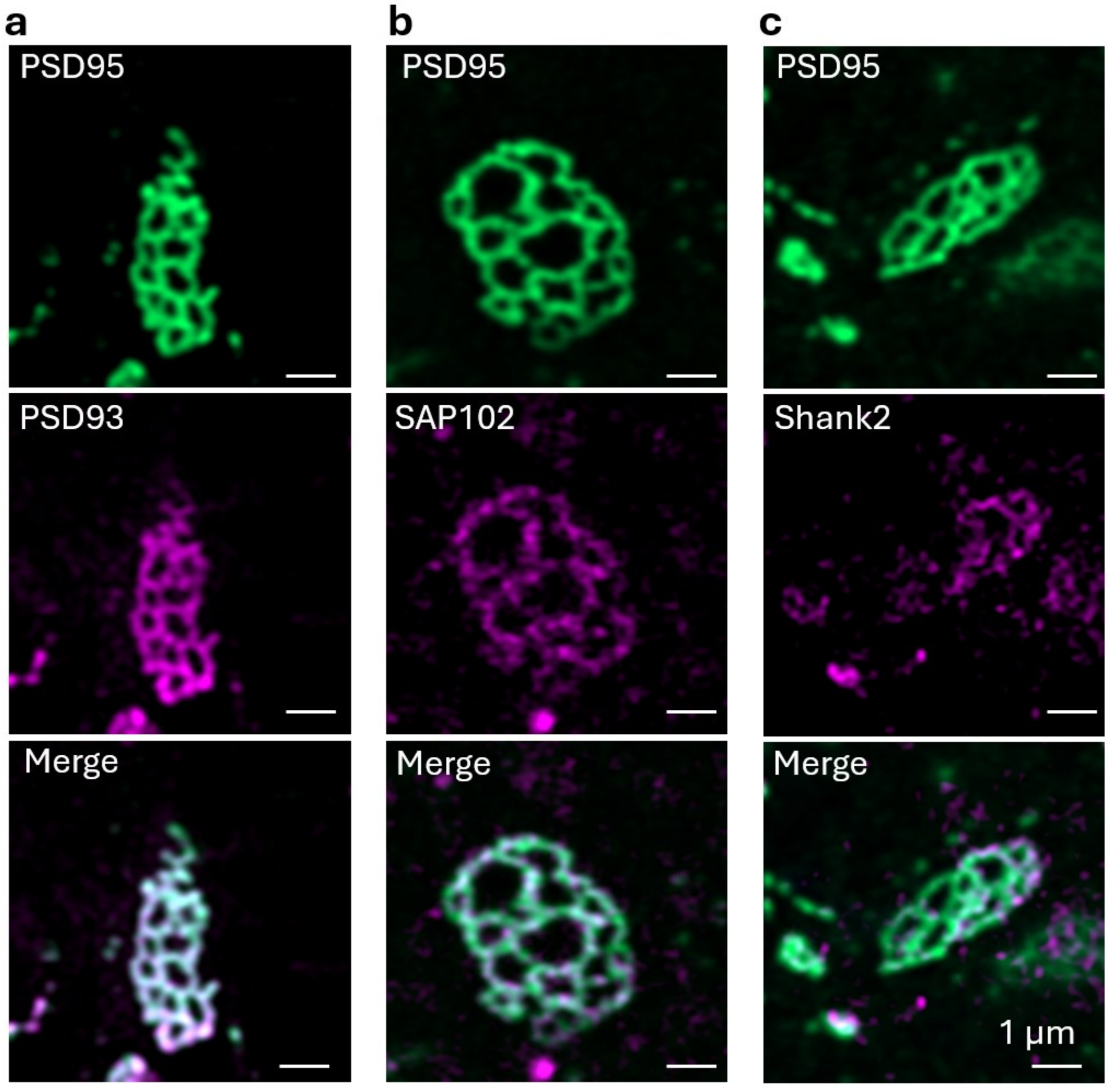


***Supplementary Figure 1.*** *Organisation of Scaffold Proteins in Honeycomb Synapses. PSD95-eGFP Honeycomb Synapses were visualised, alongside PSD93 (a), SAP102 (b) and Shank2 (c), showing the separate images, and the merge composite.*

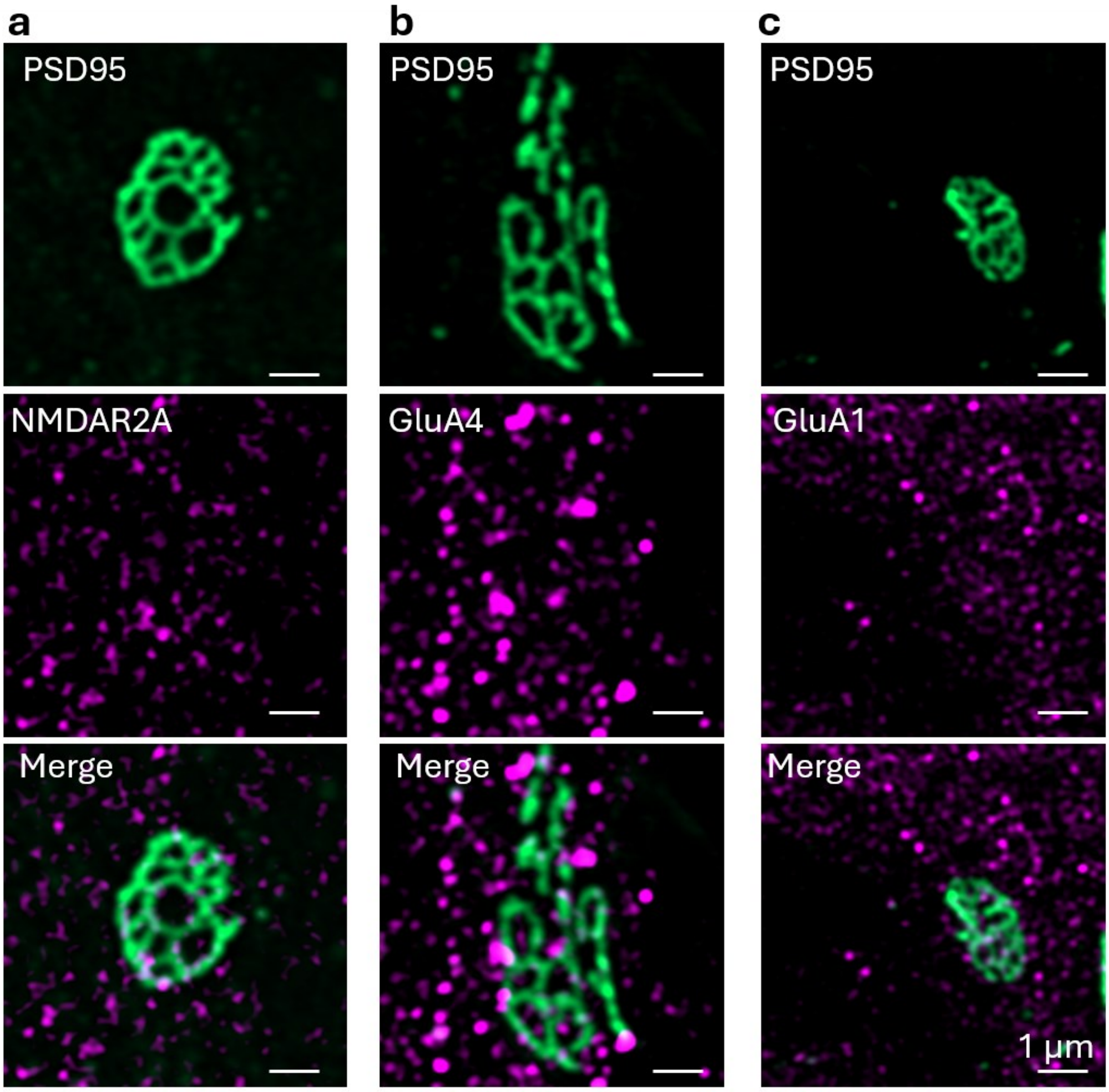


***Supplementary Figure 2. O****rganisation of Glutamatergic Receptors in Honeycomb Synapses. PSD95-eGFP Honeycomb Synapses were visualised, alongside NMDAR2A (a), GluA4 (b) and GluA1 (c), showing the separate images, and the merge composite.*

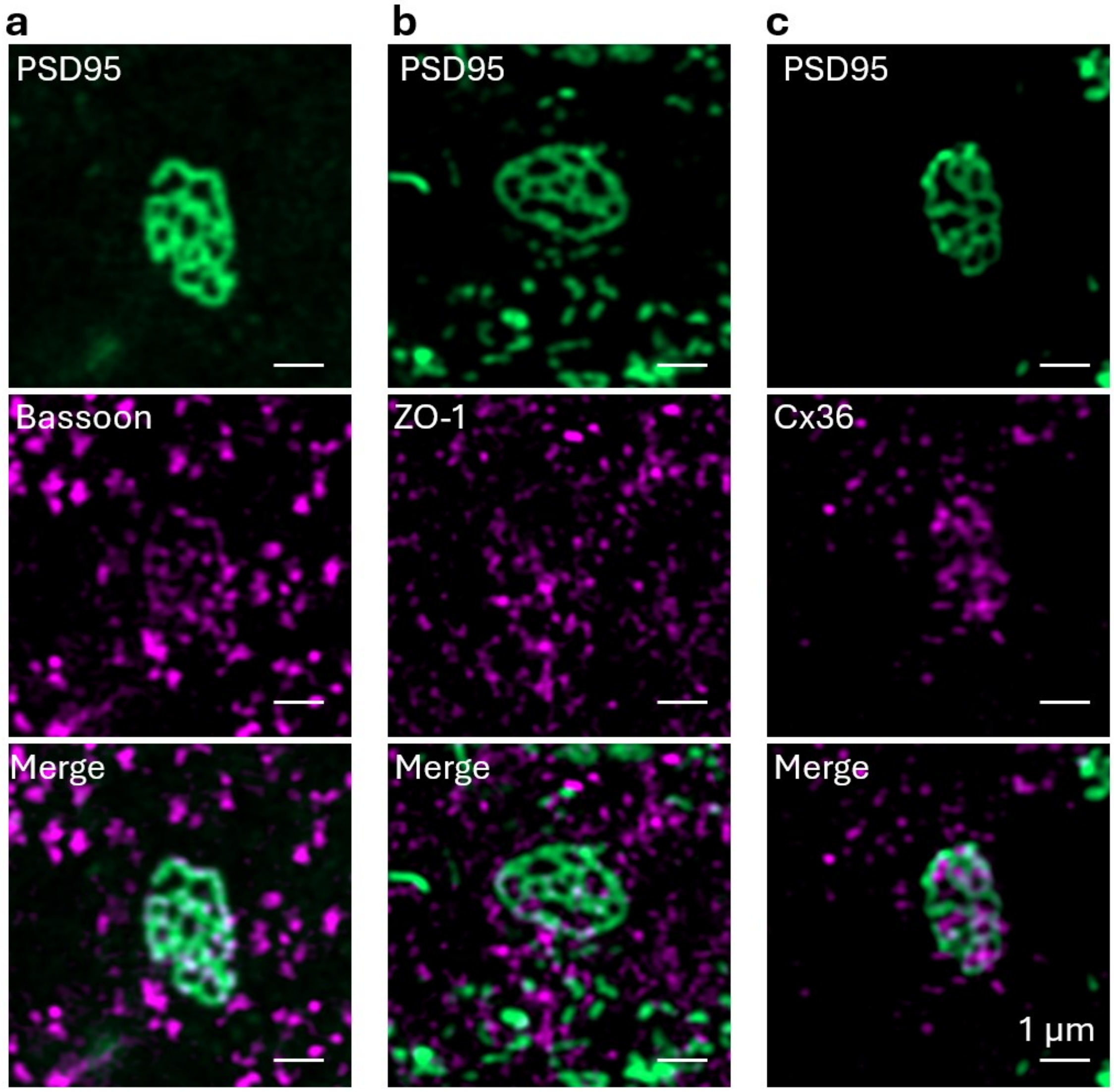


***Supplementary Figure 3.*** *Organisation of Vesicular Release and Gap Junction Proteins in Honeycomb Synapses. PSD95-eGFP Honeycomb Synapses were visualised, alongside Bassoon (a), ZO-1 (b) and Cx36 (c), showing the separate images, and the merge composite.*

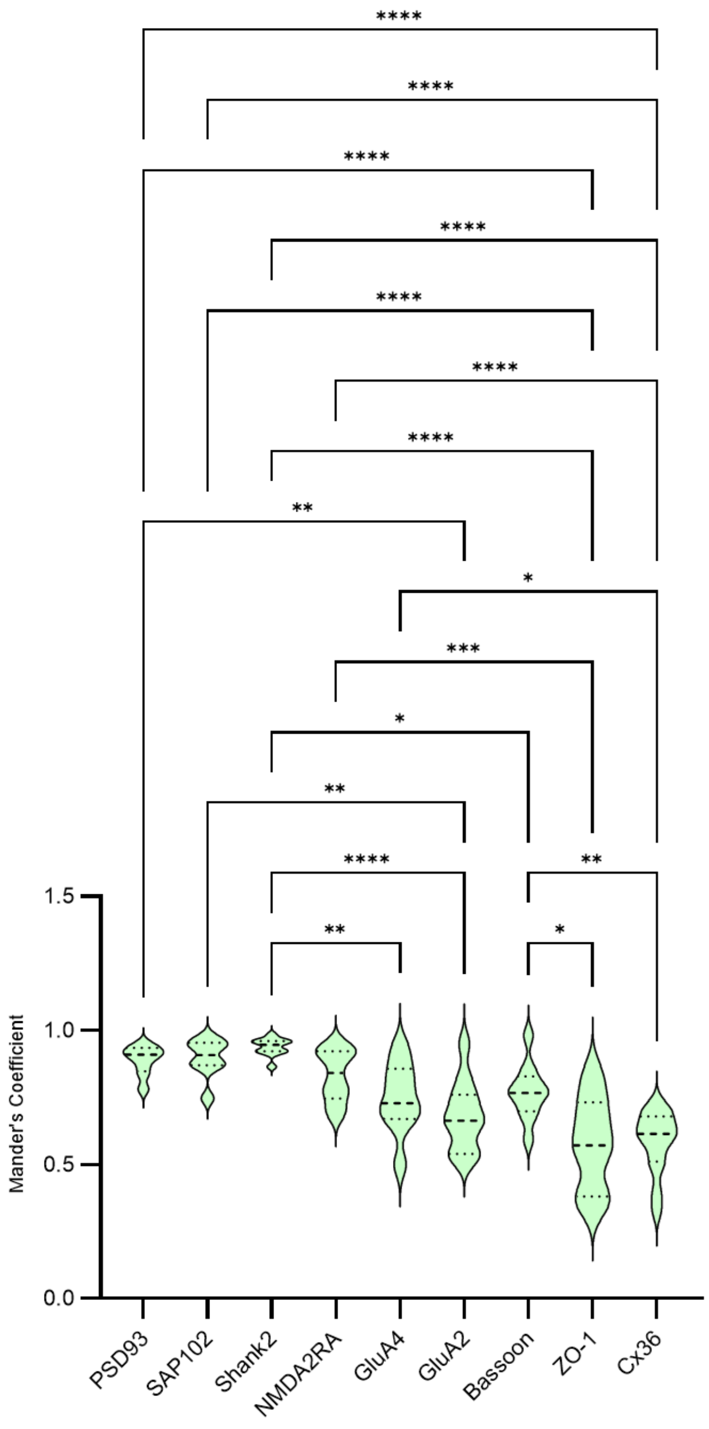


***Supplementary Figure 4.*** *Bar chart plotting the degree of colocalization (Mander's Coefficient) of each synaptic marker within the Honeycomb Synapse Matrix as discerned from PSD95-eGFP expression. * = p<0.05, ** p<0.01, *** p<0.001, **** p<0.0001*

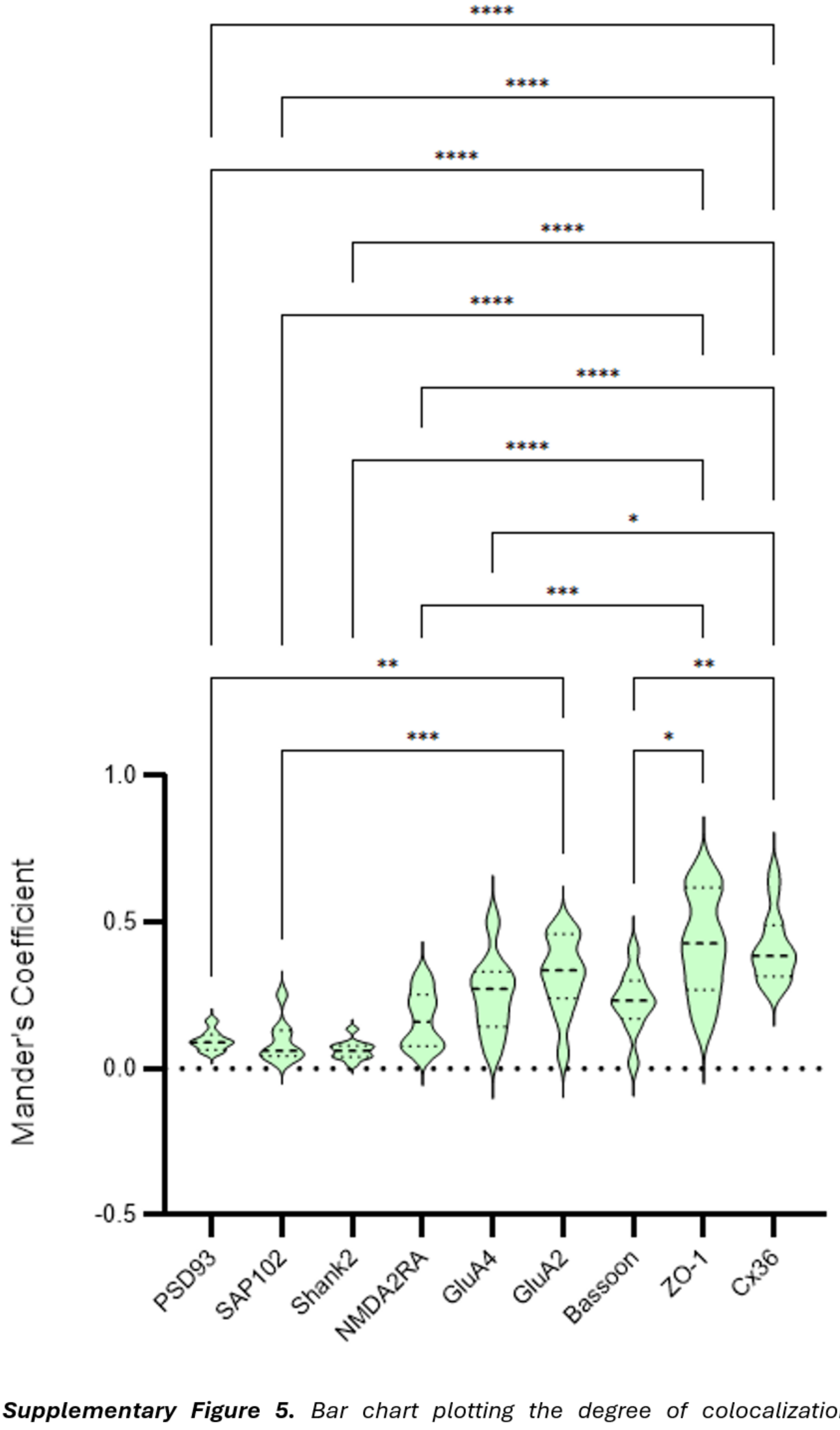


***Supplementary Figure 5.*** *Bar chart plotting the degree of colocalization (Mander's Coefficient) of each synaptic marker within the Honeycomb Synapse perforations as discerned from PSD95-eGFP expression. * = p<0.05, ** p<0.01, *** p<0.001, **** p<0.0001*

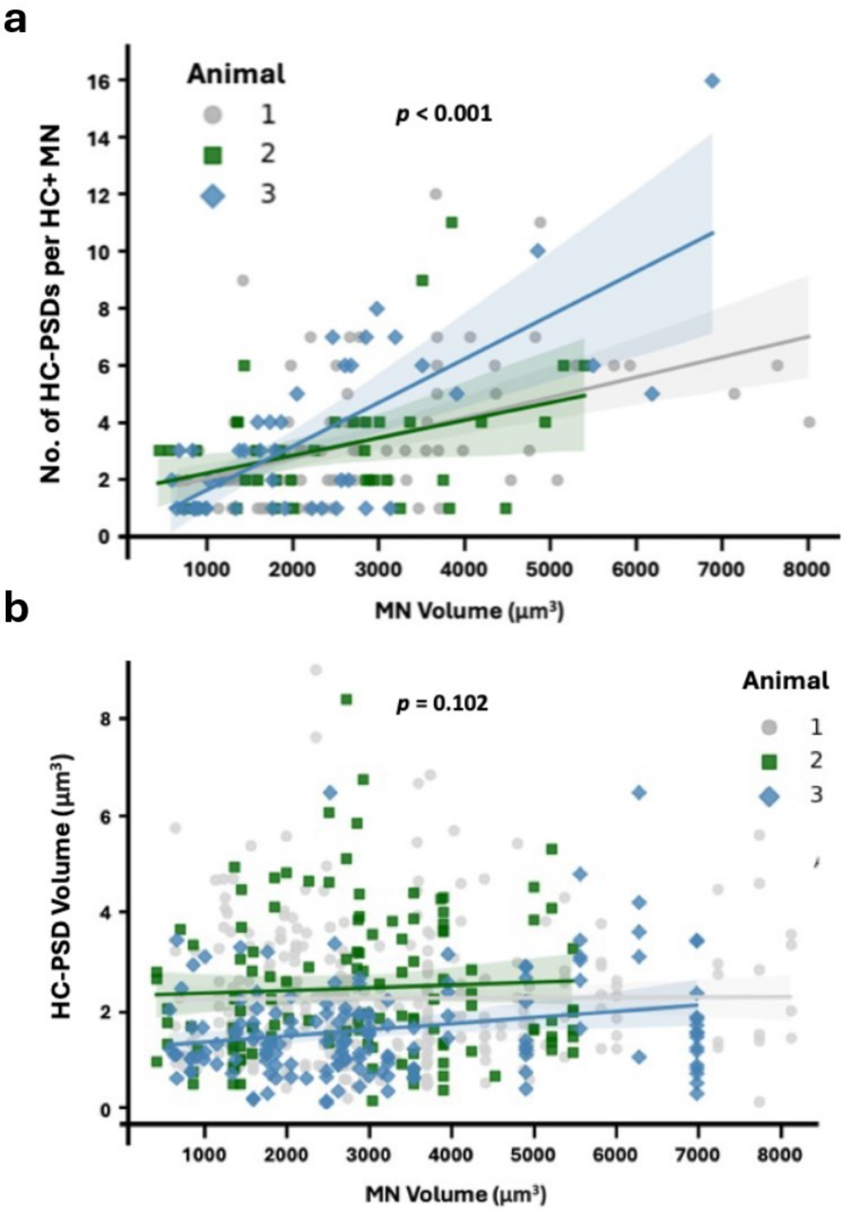


***Supplementary Figure 6.*** *a, Scatter plot showing number of colocalised Honeycomb Synapse postsynaptic densities (HC-PSD) per the volume of the motoneuron (MN) that harboured Honeycomb Synapses (HC+ MN). The data demonstrates a positive correlation between MN size and the number of Honeycomb Synapses harboured by the MN. b, Scatter plot showing HC-PSD volume as a function of MN volume by animal. The data demonstrates there is no correlation between MN size and Honeycomb Synapse size. Data was separated by each of the 3 animals as per the keys to the right of each graph.*

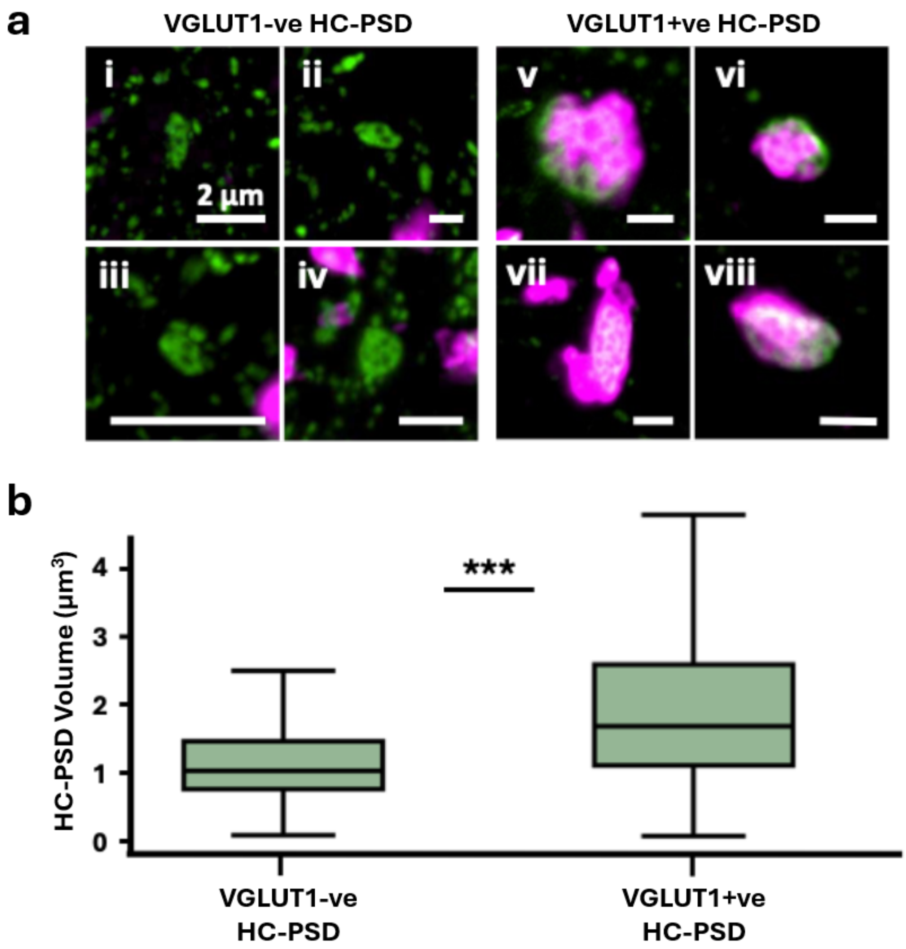


***Supplementary Figure 7.*** *a, Honeycomb Synapses were identified from PSD95 structures harbouring at least 2 perforations. Whilst most Honeycomb Synapses were observed opposed to VGLUT1 (VGLUT1+ve HC-PSD; examples i-iv), a small portion were not (VGLUT1-ve HC-PSD; examples v-viii). b, Graph plotting the size of VGLUT1 –ve and VGLUT1 +ve HC-PSDs. VGLUT1 +ve HC-PSDs are significantly larger than VGLUT1 –ve HC-PSDs ($\beta$ = 0.746, $p < 0.001$) .*
